# iPSC-Derived PSEN2 (N141I) Astrocytes and Microglia Exhibit a Primed Inflammatory Phenotype

**DOI:** 10.1101/2022.12.05.518134

**Authors:** Michael A. Sullivan, Samuel D. Lane, Sarah R. Ball, Margaret Sunde, G. Gregory. Neely, Cesar Moreno, Eryn L. Werry, Michael Kassiou

## Abstract

**Background:** Widescale evidence points to the involvement of glia and immune pathways in the progression of Alzheimer’s disease (AD). AD-associated iPSC-derived glial cells show a diverse range of AD-related phenotypic states encompassing cytokine/chemokine release, phagocytosis and morphological profiles, but to date studies are limited to cells derived from PSEN1, APOE and APP mutations or sporadic patients. The aim of the current study was to successfully differentiate iPSC-derived microglia and astrocytes from patients harbouring an AD-causative PSEN2 (N141I) mutation and characterise the inflammatory and morphological profile of these cells.

**Methods:** iPSCs from three healthy control individuals and three familial AD patients harbouring a heterozygous PSEN2 (N141I) mutation were used to derive astrocytes and microglia-like cells and cell identity and morphology were characterised through immunofluorescent microscopy. Cellular characterisation involved the stimulation of these cells by LPS and Aβ_42_ and analysis of cytokine/chemokine release was conducted through ELISAs and multi-cytokine arrays. The phagocytic capacity of these cells was then indexed by the uptake of fluorescently labelled fibrillar Aβ_42_.

**Results:** AD-derived astrocytes and microglia-like cells exhibited an atrophied and less complex morphological appearance than healthy controls. AD-derived astrocytes showed increased basal expression of GFAP, S100β and increased secretion and phagocytosis of Aβ_42_ while AD-derived microglia-like cells showed decreased IL-8 secretion compared to healthy controls. Upon immunological challenge AD-derived astrocytes and microglia-like cells show exaggerated secretion of the pro-inflammatory IL-6, CXCL1, ICAM-1 and IL-8 from astrocytes and IL-18 and MIF from microglia.

**Conclusion:** Our study showed, for the first time, the differentiation and characterisation of iPSC-derived astrocytes and microglia-like cells harbouring a PSEN2 (N141I) mutation. PSEN2 (N141I)-mutant astrocytes and microglia-like cells presented with a ‘primed’ phenotype characterised by reduced morphological complexity, exaggerated pro-inflammatory cytokine secretion and altered Aβ_42_ production and phagocytosis.

## Introduction

Alzheimer’s disease (AD) is currently a largely unmet worldwide clinical burden, despite being the subject of many clinical trials over the last decade. AD occurs either as early onset familial AD (fAD) around the fourth decade of life or late stage sporadic AD (sAD) which usually develops past 65 years old. sAD is the more common form of the disease (> 95 % frequency) with the biggest genetic risk factors for developing sAD including triggering receptor expressed on myeloid cells 2 (TREM2) and the ε4 allele of the Apolipoprotein E (APOE) gene (1). fAD accounts for less than 5 % of total AD cases and the causative mutations are inherited in an autosomal dominant fashion, with over 200 different mutations occurring in the amyloid precursor protein (APP), presenilin-1 (PSEN1) and presenilin-2 (PSEN2) genes (1, 2). PSEN1/2 subunits are at the catalytic core of the γ-secretase complex and as such, mutations in these subunits are central to the amyloidogenic processing of APP within fAD (3, 4). Of all the fAD-causing mutations in PSEN2 that have been identified, the N141I missense mutation is the most prevalent AD-causing PSEN2 mutation and has a very high clinical penetrance (> 95 %) (5). The etiology of AD is not well understood but both the sporadic and familial forms of the disease have common neuropathological features which most noticeably include gross brain atrophy, neuroinflammation, insoluble parenchymal amyloid-β (Aβ) deposits and intracellular neurofibrillary tangles containing hyperphosphorylated tau (6).

Increasing evidence suggests glial cells are central to disease-modifying dysfunctions in AD pathogenesis. Genome-wide association studies have identified several genetic risk loci that implicate the innate immune system in the development of AD (7). Within the brain of AD patients, increased microglial activation is observed during the prodromal and potentially pre-clinical stages of AD and is present in both mildly and severely cognitively impaired individuals (8, 9). Additionally, astrocytes in the post-mortem brain of AD patients exhibit significant cellular atrophy, upregulate the cytoskeletal protein glial fibrillary acidic protein (GFAP) and internalize Aβ (10-13). While post-mortem studies find microglia and astrocytes surrounding Aβ aggregates in high numbers, this research fails to answer when and how microglia and astrocytes specifically respond to Aβ, and whether this is beneficial or detrimental for AD progression (14-16). Furthermore, preclinical transgenic mouse models of AD are limited in their ability to imitate the early stages of the pathological cascade, show significant species differences to humans and show poor clinical translation of therapeutics (17). As such, there remains a clear need to investigate cell-specific and cell-autonomous changes occurring early in disease progression, by examining human glial cells *in vitro*.

Induced pluripotent stem cell (iPSC)-derived CNS cells closely mimic their *in vivo* counterparts and provide a new opportunity to characterise AD-associated functions within these cells *in vitro* (18, 19). Previous publications have reported the successful *in vitro* differentiation of iPSCs to microglia-like cells and astrocytes with high yield and purity (20-22). iPSC-derived astrocytes and microglia-like cells from both sAD and fAD origin have been utilised to investigate a variety of AD-associated changes in morphology, cytokine/chemokine release and phagocytosis. The existing body of research suggests that iPSC-derived glial cells show complex behaviours, likely indicative of a diverse range of AD-related phenotypic states encompassing cytokine/chemokine release, phagocytosis and morphological profiles. APP and PSEN1 mutant fAD-derived astrocytes and sAD-derived microglia-like cells exhibit a general increase in pro-inflammatory cytokine production both basally and in response to inflammatory stimulation such as IL-6, IL-8 and tumor necrosis factor-α (TNF-α). However, there was variability in profiles of astrocyte chemoattractant secretion and microglial pro-inflammatory cytokine release between the different AD patient origins (23-29).

The existing body of research examining the molecular and cellular behaviour of iPSC-derived glia has focused on iPSCs established from APP, PSEN1 or sAD donors, with no studies having directly examined astrocytes/microglia derived from patients carrying PSEN2 mutations. Given the lack of research on iPSC-derived PSEN2 mutant glial cells, it is important to establish the AD-associated cellular characteristics of PSEN2-mutant astrocytes and microglia. This will provide further understanding of common inflammatory or other glial-specific mechanisms which may help guide drug discovery towards a pan-AD treatment. The aim of the current study was to successfully differentiate iPSC-derived microglia and astrocytes from patients harbouring a PSEN2 (N141I) mutation and investigate the morphological, inflammatory and phagocytic phenotype presented.

## Materials and Methods

### iPSC Lines

Human control iPSCs were obtained from the Cedars-Sinai iPSC Core cell repository (Los Angeles, USA). Heterozygous fAD PSEN2 (N141I) iPSCs were obtained from both the Cedars-Sinai (Los Angeles, USA) and New York Stem-Cell Foundation cell repositories (New York, USA). All lines were generated from dermal fibroblasts obtained from skin punch biopsies, reprogrammed using either a non-integrating episomal plasmid or mRNA transfection and showed normal karyotyping. A summary of the iPSC line characteristics is shown in Table 1.

**Table 1.**
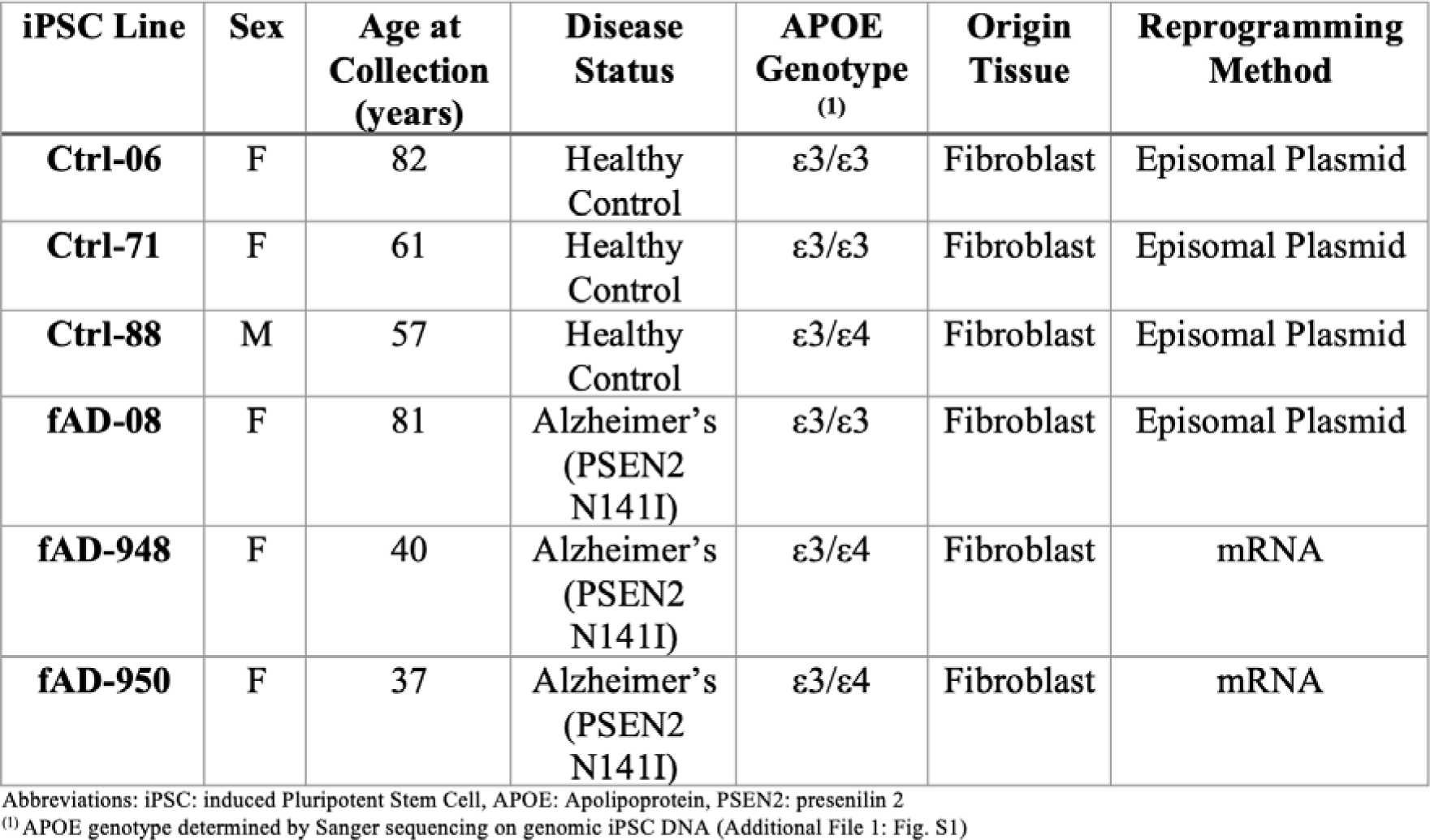
Summary of iPSC line characteristics

### iPSC Culture

iPSCs were cultured in feeder-free conditions on Matrigel-coated (0.08 mg/well) 6-well tissue culture plates in mTesR1 media (StemCell Technologies) at 37 °C and 5 % CO_2_. Cultures were fed daily. For passaging of iPSC cultures, differentiated colonies were manually scratched off the bottom of the well and media was aspirated. Fresh media was added and passaged 1:6 using the StemPro EZpassage tool (Life Technologies) as per the manufacturer’s instructions.

### NPC Derivation

Derivation of neural progenitor cells (NPCs) from iPSCs closely followed a previously published protocol (22). iPSCs were cultured with the addition of 10 ng/mL StemBeads fibroblast growth factor 2 (FGF-2) (StemCultures). Upon reaching 80 % confluency, the media was changed to TeSR-E8 supplemented with 10 ng/mL StemBeads FGF-2 and 10 _μ_M Y-27632 (Sigma-Aldrich). After 24 hrs, cells were washed with phosphate buffered saline (PBS), dissociated with gentle cell dissociation reagent (StemCell Technologies) and incubated for 10 min (37 °C, 5 % CO_2_). Cells were gently pipetted up and down to ensure all cells had dislodged and formed a single cell suspension. Cells were then centrifuged (300 *g*, 5 min) and plated at a density of 90,000 cells/mL in STEMdiff Neural Induction Medium (StemCell Technologies) with SMADi supplement (StemCell Technologies) (NIM) and 10 μM Y-27632 on an ultra-low attachment U-bottom 96-well plate (100 μL/well) (Costar) and incubated at 37 °C, 5 % CO_2_. Three-quarters of the media was changed daily for 4 days, ensuring the embryoid bodies were not removed. Embryoid bodies were aspirated using a 200 μL wide-bore pipette tip and plated onto Matrigel-coated (0.08 mg/well) 6-well plates (10-13 embryoid bodies/well) in NIM. Media was fully replaced daily for 7 days. Cells were washed with Dulbecco’s Modified Eagle Medium (DMEM/F12) and incubated for 70 min (37°C, 5 % CO_2_) in STEMdiff Neural Rosette Selection Reagent (StemCell Technologies). Rosettes were lifted by gently dispensing DMEM/F12 onto the colonies, centrifuged (350 *g*, 5 min), plated onto a Matrigel-coated (0.08 mg/well) 6-well plate in NIM and incubated for 24 h (37 °C, 5 % CO_2_). Media was switched to neural progenitor cell media containing DMEM/F12, 1x N2 (Invitrogen), 1x B27-RA (Invitrogen) and 20 ng/ml FGF_2_ (Abcam) and changed daily. NPCs were passaged after 4 days using accutase (Sigma-Aldrich) (5 min, 37°C, 5 % CO_2_) and maintained on Matrigel-coated (0.08 mg/well) 6-well plates. Further passaging was done approximately 1:3 every week.

### Astrocyte Derivation

Differentiation of NPCs into astrocytes followed a previously published protocol (22). NPCs were dissociated using accutase (5 min, 37 °C, 5 % CO_2_), centrifuged (300 *g*, 5 min), plated at 15,000 cells/cm^2^ on Matrigel-coated (0.08 mg/well) 6-well plates in NPC medium and incubated for 24 h (37 °C, 5 % CO_2_). Media was switched to astrocyte medium (astrocyte basal medium (ScienCell), 2 % fetal bovine serum, astrocyte growth supplement and 10 U/mL penicillin/streptomycin solution) and were fed every other day. Cells were passaged with accutase (5 min, 37 °C, 5 % CO2) when reaching 90 % confluency, centrifuged (300 *g*, 5 min) and plated at the original plating density. After 30 days in astrocyte medium, astrocyte identity was then validated using immunofluorescence and used for experiments.

### Microglia Derivation

Differentiation of iPSCs to hematopoietic progenitor cells (HPCs) was done using Stem Cell Technologies STEMdiff Hematopoietic Kit (StemCell Technologies). Upon reaching 80 % confluency, iPSCs were detached using gentle cell dissociation reagent (StemCell Technologies) (10 min, 37 °C, 5 % CO_2_) and plated at 40 colonies/well on Matrigel-coated (0.08 mg/well) 12-well plates in mTeSR1 media (StemCell Technologies) supplemented with mTeSR1 5× Supplement (StemCell Technologies), ensuring the colony sizes were around 50-100 μm in diameter. The following day mTeSR1 medium was replaced with media A (Hematopoietic basal medium with supplement A at 1:200). After 48 h, a half media change was conducted with media A. Twenty-four hours later, media was switched to media B (Hematopoietic Basal Medium with supplement B at 1:200). A half media change was conducted 2, 4 and 7 days later, ensuring not to disturb the floating cell population. After 9 days in media B, media was completely removed and washed once with DMEM/F12 to ensure all non-adherent cells were collected. These cells were hematopoietic progenitor cells. The cell-containing supernatant was centrifuged (300 *g*, 5 min) and plated at 200,000 cells/well on Matrigel-coated (0.08 mg/well) 6-well plates containing 2 mL STEMdiff Microglia Differentiation Media (Microglia Basal Medium with supplement 1 at 1:10 and supplement 2 at 1:250). STEMdiff Microglia Differentiation Media (1 mL) was added every other day. After 12 days both the semi-adherent and non-adherent cells were collected, centrifuged (300 *g*, 5 min), resuspended in a 1:1 mixture of cell supernatant and fresh STEMdiff Microglia Differentiation Media and plated on a Matrigel-coated (0.08 mg/well) 6-well plate. STEMdiff Microglia Differentiation Media (1 mL) was added every other day for a further 12 days. Non-adherent cells were collected and adherent cells were detached using accutase (5 min, 37 °C, 5 % CO_2_), centrifuged (300 *g*, 5 min) and plated on Matrigel-coated surfaces (depending on application) at 100,000 cells/cm^2^ in STEMdiff Microglia Maturation Media (STEMdiff Microglia Differentiation Media with supplement 3 at 1:250). Half of the initial plating media volume of STEMdiff Microglia Maturation Media was added every other day for 8 days, at which point the microglia-like cells were used for analysis.

### Immunofluorescence

NPCs and iPSC-derived astrocytes were plated at 45,000 cells/well and iPSCs were plated as colonies on Matrigel (0.08 mg/mL)-coated 8-well glass chamber slides. 24 h post plating, cell media was aspirated and cells were washed 3x with PBS. Cells were then fixed with 4 % paraformaldehyde (10 min, 22 °C), washed 3x with PBS and permeabilised with 0.1 % TritonX-100 (10 min, 22 °C). At day 24 of the microglia differentiation, iPSC-derived microglia were plated on Matrigel-coated 8-well glass chamber slides at 100,000 cells/well in STEMdiff Microglia Maturation Media. After microglia maturation for 8 days, media was removed without washing then cells fixed with 4 % paraformaldehyde (10 min, 22 °C), washed 3x with PBS and permeabilised with 0.1 % TritonX-100 (10 min, 22 °C). Blocking for all cell types was done in 5 % (v/v) fetal bovine serum in PBS (blocking solution; 1 h, 22°C). Primary antibodies were diluted in blocking solution and incubated overnight (4 °C). The following day, cells were washed 3x with PBS and incubated with the appropriate Alexa 488/594-conjugated secondary antibody in blocking solution (1 h, 22 °C) (antibody details and dilutions are shown in Additional file 1: Tables S1 and S2). Cells were washed 3x in PBS and mounted on a glass coverslip in fluroshield with DAPI (Sigma-Aldrich). Confocal microscopy was performed using the LSM800 (Zeiss, ZEN Blue software) and images processed using FIJI image analysis software.

### Astrocyte Morphology and Fluorescent Intensity Analysis

After cells were fixed and stained as above, imaging and analysis followed Jones *et al*., (23). To ensure consistent imaging between samples, the mono-directional scan speed, laser power, digital gain, offset and pinhole (set to 1 airy unit) were kept constant for all experiments. Images were taken as 34 z-sections spaced at 0.47 μm intervals on a 10x air objective and processed as maximum intensity projections in FIJI. A minimum of 80 cells were imaged from 2 random fields for each cell line. Morphological analysis of astrocytes was carried out by visually binning each cell into one of three categories defined by Jones *et al*., (23): arborized (defined as having greater than 2 distinct processes where the longest extended further than the width of the cell body), bipolar (defined as having 2 distinct processes where the longest extended further than the width of the cell body) and process devoid (defined as having the longest process extended less than the width of the cell body). Cell perimeter, area, GFAP/S100β fluorescent intensity and circularity was calculated using FIJI. Circularity was defined as 4*π*Area/Perimeter^2^, with a value of 1 describing a perfectly circular object. Cell volume was calculated using the z-stack described above and calculated using the ‘3D object counter’ plugin within FIJI.

### Microglia Morphology Analysis

iPSC-derived microglia-like cells were fixed and stained for the marker IBA1 and confocal microscopy was performed using the LSM800 (Zeiss, ZEN Blue software). Image processing and morphological quantification was conducted in FIJI and followed the protocol previously developed by Young and Morrison (2018). IBA1 images were converted to grayscale and edge features were enhanced by applying an unsharp mask (pixel radius: 3 and mask weight: 0.6). Individual pixel background noise was removed through the ‘despeckle’ function. Images were then manually thresholded to a value that included processes without background signal to create a binary image. Background noise was then removed using the ‘despeckle’ function and pixels less than 2 pixels apart joined using the ‘close’ function. Bright pixel outliers were removed through the ‘remove outliers’ function (pixel radius: 2 and threshold: 50). The process binary images were then automatically skeletonised and analysed using the ‘analyse skeleton’ function. Measurements that contained < 2 branches were background noise and removed from the analysis.

### Aβ_42_ Production and Purification

Overexpression and purification of unlabeled Aβ_42_ was performed according to Walsh *et al*., (30). Briefly, large cultures of Escherichia coli BL21 Gold (DE3) were incubated at 37 °C with shaking, induced with 0.5 mM isopropyl β-D-1-thiogalactopyranoside and harvested by centrifugation. Purification involved a series of sonication and centrifugation steps followed by resuspension of inclusion bodies in 8 M urea. Anion exchange chromatography using diethylaminoethyl cellulose beads was performed and protein eluted with 150 mM NaCl. Aβ_42_ elution was assessed by electrophoresis, using by Novex 10–20% Tricine SDS-PAGE gels. Fractions containing high levels of protein were combined and aliquoted, and lyophilised. A single aliquot was resuspended in 6 M GuHCl 20 mM NaH_2_PO_4_ pH 8.0 and filtered through a 30 kDa molecular mass cut-off centrifugal filtration device (Amicon Ultra-15) and kept at room temp for 1 h. A Sepahedex G-25 column was equilibrated with 50 mM ammonium acetate pH 8.5 and protein sample loaded onto the beads. Fractions were collected in Protein LoBind Eppendorf and immediately placed on ice. Aβ_42_ elution was assessed by electrophoresis, using by Novex 10–20% Tricine SDS-PAGE gels and aliquots were snap-frozen in liquid nitrogen and lyophilised. Freeze dried protein was resuspended in Hexafluoro-2-propanol at 1 mg/ml and incubated for 30 min at RT. Protein was snap-frozen, lyophilised, and stored at –80 °C until use. Lyophilized protein was gently resuspended in 10 mM NaOH at 2 mg/mL and bath sonicated in ice water for 1 min. 20 mM NaH_2_PO_4_ pH 6.6 was added at 1:2 ratio v/v (NaOH: NaH_2_PO_4_). The sample was centrifuged at 16000 × *g* for 10 min at 4 °C. Concentration was measured using ε275 = 1450 M^-1^cm^-1^ and protein was maintained on ice until use.

### IL-6 Cytokine ELISA

iPSC derived astrocytes were plated on Matrigel-coated (0.08 mg/well) 96-well plates at a density of 40,000 cells/well in astrocyte media. 24 h after plating, media was aspirated and cells were exposed to 10 or 50 μg/mL lipopolysaccharide (LPS) (*E. coli* strain 0111:B4; Sigma-Aldrich) (MilliQ H_2_O vehicle) or to 5 or 10 μM Aβ_42_ in astrocyte media for 24 h. Media was removed, centrifuged (10,000 *g,* 1 min) and the supernatant stored at -80 °C. IL-6 ELISA was performed according to the manufacturer’s protocol (R&D Systems product number DY206), with slight amendments including: capture antibody used at 2 μg/mL, detection antibody used at 50 ng/mL and standard range used from 0.586-600 pg/mL. Absorbance at 450 nm was recorded using a BMG POLARstar Omega and analysed using a sigmoidal dose-response (variable slope) model, with unknowns interpolated in GraphPad Prism 9.

### PCR and Sanger Sequencing

iPSCs were grown to 80 % confluency on Matrigel-coated 6-well plates. Genomic DNA was extracted using the Purelink Genomic DNA Mini Kit (Invitrogen), and concentration and purity was determined using the Nanodrop ND-1000 Spectrophotometer. The APOE region was amplified using PCR, where 500 ng DNA was mixed with 25 uL NEBNext High-Fidelity 2X PCR Master Mix, 500 nM forward primer (TCTTGGGTCTCTCTGGCTCA; Integrated DNA Technologies), and 500 nM reverse primer (GCTGCCCATCTCCTCCATC; Integrated DNA Technologies). The reaction was amplified using a T100 thermal cycler (Bio-Rad), with the following program: Initial denaturation at 98 °C for 30 s, followed by 40 cycles of denaturation for 10 s at 98 °C, annealing for 15 s at 69 °C and extension for 30 s at 72 °C ; then a final extension at 72 °C for 2 min. Electrophoresis was performed on an E-gel Powerbase V4 (Invitrogen) at 100 V for 30 min, using a 1.2% agarose E-Gel with SYBR Safe (Invitrogen), then visualised using an E-Gel Safe Imager Real Time Transilluminator (Invitrogen) to ensure specific amplification. The product band was excised from the gel, and purified using the QIAquick Gel Extraction Kit (Qiagen). The purified PCR product was then submitted to AGRF for Sanger sequencing, and sequences at SNP locations rs429358 and rs7412 were analysed using SnapGene version 6.1.1.

### Multi-Cytokine Arrays

iPSC-derived astrocytes were plated on Matrigel-coated (0.08 mg/well) 12-well plates at a density of 407,000 cells/well in 1.1 mL/well astrocyte media. 48 h after plating, cell supernatants were removed, centrifuged (10,000 *g,* 1 min) and the resulting supernatants stored at -80 °C until analysed. At day 24 of the microglia differentiation, the cells were plated on Matrigel-coated 8-well glass chambers at 100,000 cells/well in STEMdiff Microglia Maturation Media (as per section 3.2.5). After microglial maturation for 8 days, media was removed, centrifuged (10,000 *g,* 1 min) and the supernatant stored at -80 °C until analysed. In addition to basal conditions, 10 μM monomeric Aβ was added to the cells 24 h prior to the collection of media in both conditions. Protein levels were measured using the Proteome Profiler Human Cytokine Array Kit (R&D Systems product number ARY005B) as per the manufacturer’s instruction, using 1 mL and 0.5 mL of astrocyte and microglia supernatant, respectively. Each membrane was imaged using the Chemi-Doc MP system (Bio-rad) at a constant exposure time for all experiments. Signal intensity for each protein was conducted in duplicate and determined through total pixel intensity using FIJI.

### Aβ_42:40_ ELISA and BCA

iPSC derived astrocytes were plated in Matrigel-coated (0.08 mg/well) 96-well plates at a density of 40,000 cells/well in astrocyte media. Conditioned media was collected 72 h after plating and Aβ levels quantified using a human/rat β-amyloid 40 ELISA Kit and-β amyloid 42 ELISA Kit high sensitive (Wako) according to the manufacturer’s protocol. The Aβ_40_ and Aβ_42_ standard range ranged from 1-100 pmol/L and 1-20 pmol/L, respectively. For determination of total protein concentration, astrocytes were washed with PBS, lifted using accutase (5 min, 37 °C, 5 % CO_2_) and centrifuged (300 *g*, 5 min). Cells were then resuspended in RIPA buffer (Thermo Fisher Scientific) containing 1x protease inhibitor (Merck). Protein concentration per well was determined by the Pierce^TM^ BCA protein assay (Thermo Fisher Scientific) according to the manufacturer’s protocol. Within the BCA assay, the bovine serum albumin protein standard ranged from 25-2000 μg/mL. Absorbance at 450 nm was recorded using a BMG POLARstar Omega and analysed using a sigmoidal dose-response (variable-slope) model, with unknowns interpolated in GraphPad Prism 9. Interpolated Aβ_42:40_ concentrations were then normalised to total protein concentration for each condition.

### Astrocyte Viability Analysis

After exposure to 5 or 10 μM monomeric Aβ_42_ for 24 h, astrocyte viability was assessed using the Live/Dead Viability/Cytotoxicity Kit (Molecular Probes) according to manufacturer’s instructions. Each condition was represented as a percentage of basal (vehicle-control).

### Fluorescent Aβ_42_ Phagocytosis Assay

iPSC-derived astrocytes were plated on Matrigel-coated (0.08 mg/mL) 8-well glass chambers at a concentration of 25,000 cells/well. Twenty-four hours after plating astrocytes and 8-days after microglial maturation, cells were exposed to 2 μg/mL Fluo-488-labeled Aβ_42_ (Anaspec) and left to phagocytose for 2 h (37 °C, 5 % CO_2_). Media was removed, cells washed 1x with PBS, fixed with 4 % paraformaldehyde (10 min, RT) then washed a further 3x. Cells were imaged on the Cytation 3 Cell Imaging Multi-Mode Reader (BioTek, Gen5 v2 software) under a 10x air objective, ensuring a constant exposure time and laser intensity across experiments.

### Statistical Analysis

All data is presented as the mean ± SD of three cell lines with n ≥ 2 independent experiments per line. All data was analysed using a one-way ANOVA with Tukey’s multiple comparison test or by an unpaired t-test with the exception of the multi-cytokine array. The multi-cytokine array was analysed using multiple unpaired t-tests with a false discovery rate set to 5 %. Cytokines that yielded an average intensity value less than 10 % of the maximum were considered not to be above background and not included in the statistical analysis. All data was analysed in GraphPad Prism and significance was shown by * p < 0.05, ** p < 0.01 and *** p < 0.001.

## Results

### AD-Derived Astrocytes and Microglia-like Cells Show No Deficits in Differentiating From iPSCs

The generation of iPSC-derived astrocytes and microglia-like cells occurred through multiple stages, with an overview of each stage represented in Fig. 1. To generate astrocytes *in vitro,* iPSCs were first aggregated into embryoid bodies then promoted to a neuroectoderm fate. The generation of a neuroectoderm lineage was evidenced through the production of neural rosettes, presenting as neural tube-like structures. These rosettes were selectively replated, and the resulting NPCs were expanded for the subsequent few passages (Fig. 1 A). The successful generation of NPCs was confirmed through a panel of immunofluorescence markers prior to further differentiation into astrocytes. The cells stained strongly for the NPC markers nestin and Pax6 while being absent for the pluripotency marker Oct3 (Fig. 1 B-C). Nestin staining was stronger in the cytoplasm while Pax6 staining was more localised to the nucleus. This pattern is consistent with that in previously reported primary and stem-cell derived NPCs (31). These results gave us confidence that the cells generated were a pure population of successfully differentiated NPCs and were suitable for further differentiation into astrocytes.

**Fig. 1.**
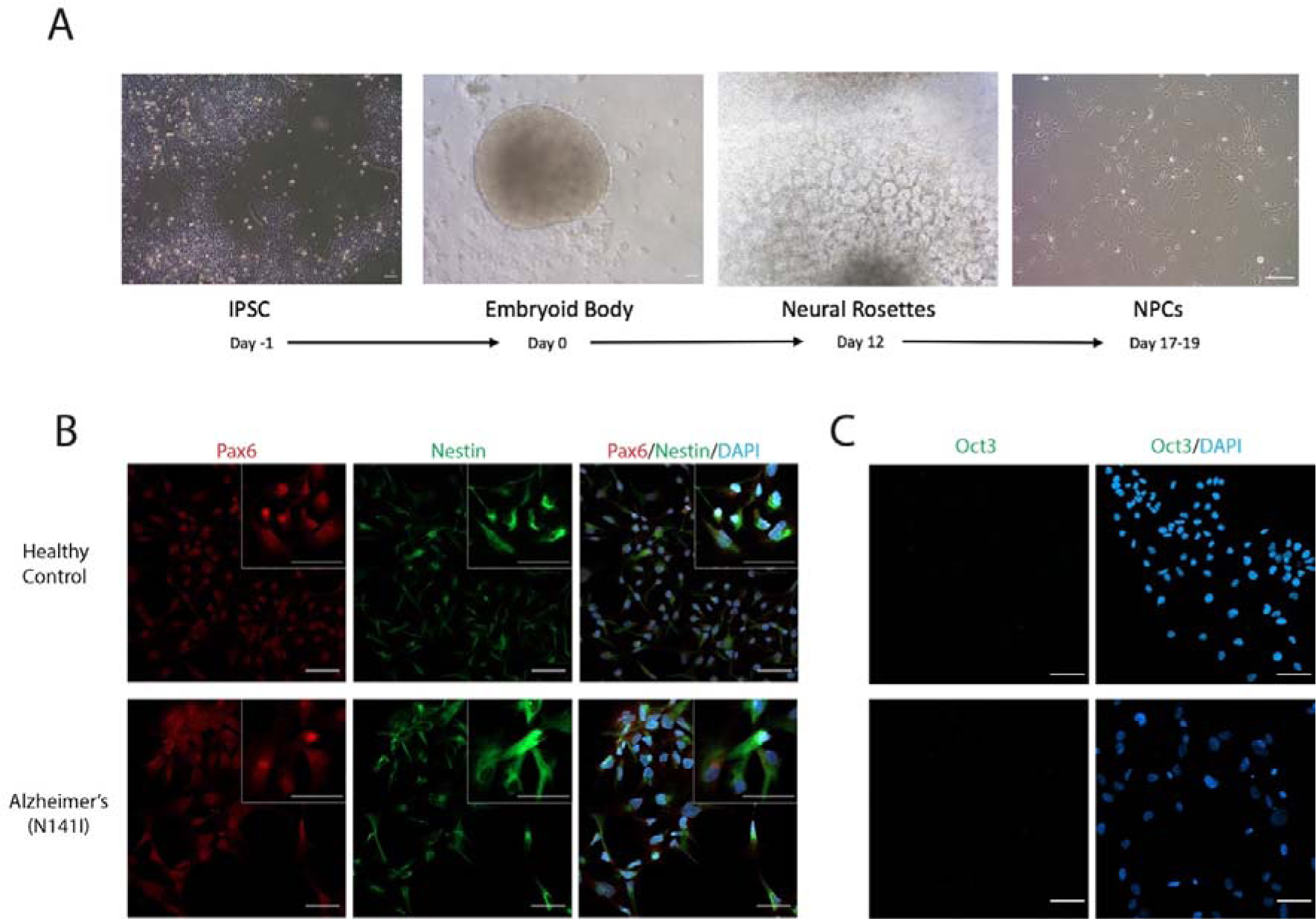
A) Overview of differentiation timeline for iPSC-derived NPCs showing representative brightfield images on the last day of each stage. Representative immunofluorescence images of iPSC-derived NPCs from healthy control lines and familial AD lines harbouring a PSEN2 (N141I) mutation. The cells were stained for B) the neural progenitor markers Pax-6 (red) and Nestin (green), C) a pluripotency marker Oct3 (green) and all nuclei were counterstained with DAPI (blue). Insert shows higher magnification. All scale bars = 50 μm. Immunofluorescence staining of all cell lines can be found in Additional File 1: Fig. S2.

After four weeks of further differentiation in astrocyte media, the resulting cells showed positive immunofluorescence staining for the canonical marker GFAP and mature astrocyte marker S100β. Furthermore, the absence of nestin staining confirmed the protocol had successfully transitioned NPCs to a pure population of mature astrocytes (Fig. 2 A and B). We found no significant difference between the Alzheimer’s PSEN2 (N141I) lines or healthy controls in the percentage of cells positive for GFAP (97 % ± 3.7 SD and 97 % ± 1.5 SD, respectively) or S100β (98 % ± 3.1 % and 98 % ± 1.9 SD, respectively) (Fig. 2 C and D).

**Fig. 2.**
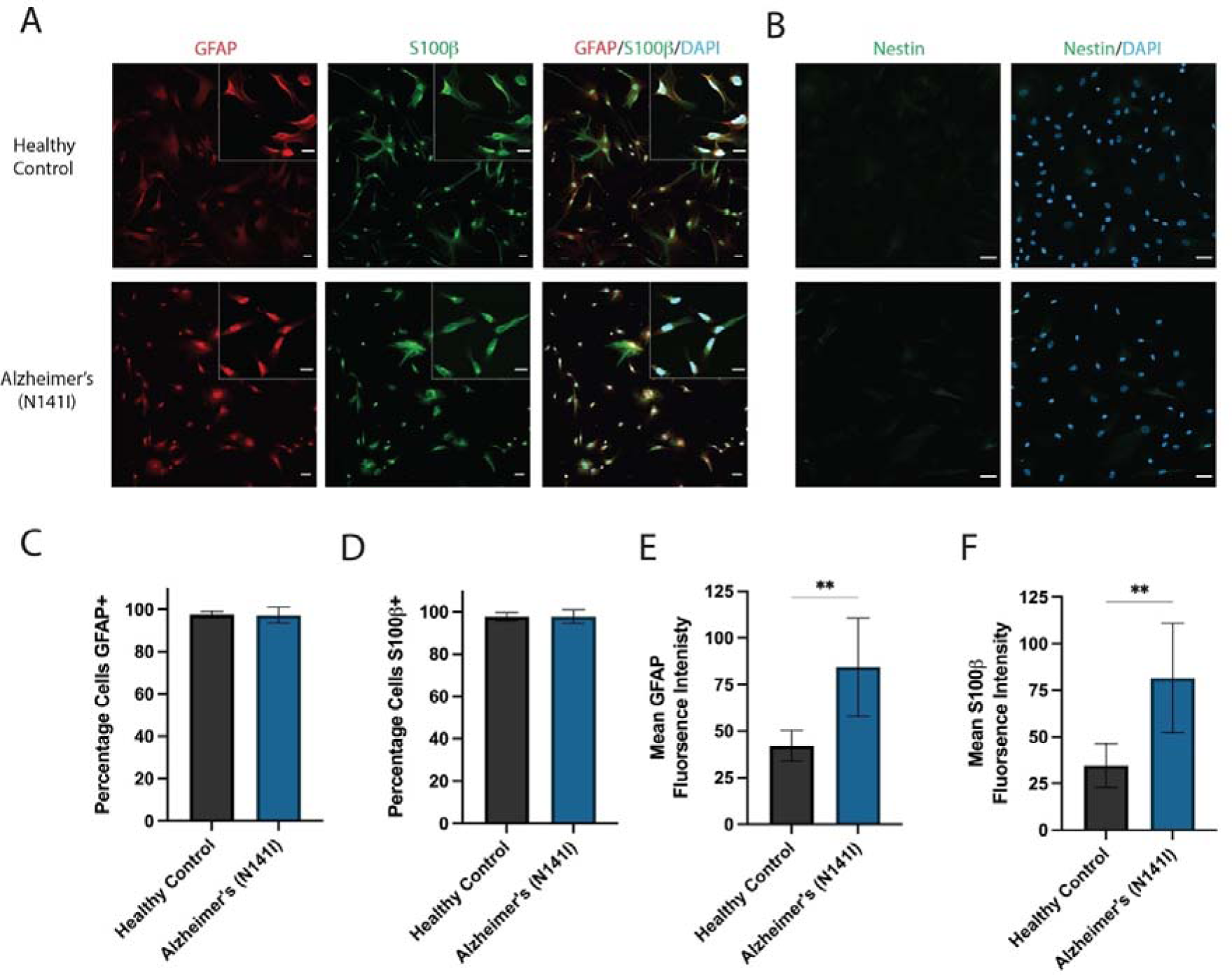
A) Representative immunofluorescence images of iPSC-derived astrocytes from healthy control lines and familial AD lines harbouring a PSEN2 (N141I) mutation. The cells were stained for astrocyte markers GFAP (red) and S100β (green) and B) the NPC marker nestin (green) and all nuclei were counterstained with DAPI (blue). Insert shows higher magnification. All scale bars = 50 μm. The percentage of iPSC-derived astrocytes positive for (C) GFAP and D) S100β. Mean fluorescence intensity of E) GFAP and F) S100β per astrocyte. An unpaired t-test was used to test whether there were statistically significant differences between the means of AD-derived and healthy control astrocytes. Each bar displays the mean ± SD of three cell lines with n ≥ 2 independent experiments per line. (** p < 0.01). Immunofluorescence staining of all cell lines can be found in Additional file 1: Fig. S3.

Intermediate filament changes are known to be present within reactive astrocytes (32), as such, we conducted fluorescent intensity analysis of GFAP and S100β to gain an initial insight into the activation profile of AD-derived astrocytes. We observed the fluorescence intensity of GFAP to be significantly elevated in AD-derived astrocytes compared to healthy controls (84.3 a.u ± 26.3 SD and 42.1 a.u ± 8.19 SD, respectively) (p < 0.01; Fig. 2 E). A significant increase in S100β intensity from AD-derived astrocytes was also observed relative to healthy controls (81.6 a.u ± 29.4 SD and 34.7 a.u ± 11.6 SD, respectively) (p < 0.01; Fig. 2 F).

To generate iPSC-derived microglia-like cells, we followed a method based on a previously published protocol which produced microglia-like cells with gene expression profiles and functional activity very similar to those of primary human fetal and adult microglia (21, 33). In short, iPSCs were plated as small colonies, patterned into floating HPC intermediates and then further differentiated into microglia-like cells (overview shown in Fig. 3 A). At the end of the derivation, the microglia-like cells exhibited strong staining for the microglial/monocyte marker IBA1 and microglial-enriched proteins TREM2 and CX3CR1 (Fig. 3 B and C). The percentage of the cell population positive for the three markers was high (> 90 %) and did not significantly differ between AD-derived and healthy control lines for IBA1 (98.4 % ± 3.0 SD and 96.6 % ± 3.5 SD, respectively), TREM2 (93.5 % ± 5.9 SD and 93.3 % ± 4.8 SD, respectively) or CX3CR1 (98.7 % ± 1.9 SD and 99.5 % ± 0.8 SD, respectively) (Fig. 3 D - F). Altered expression of IBA1 and TREM2 within microglia have shown to play a role in regulating cellular activation and inflammatory cytokine release (34, 35). Compared to healthy controls, AD-derived microglia-like cells showed a significant increase in the fluorescence intensity of IBA1 (41.7 a.u ± 16.6 SD and 72.9 a.u ± 26.7 SD, respectively) (p < 0.05; Fig. 3 G) and TREM2 (15.2 a.u ± 5.7 SD and 36.8 a.u ± 13.9 SD, respectively) (p < 0.01; Fig. 3 H).

**Fig. 3.**
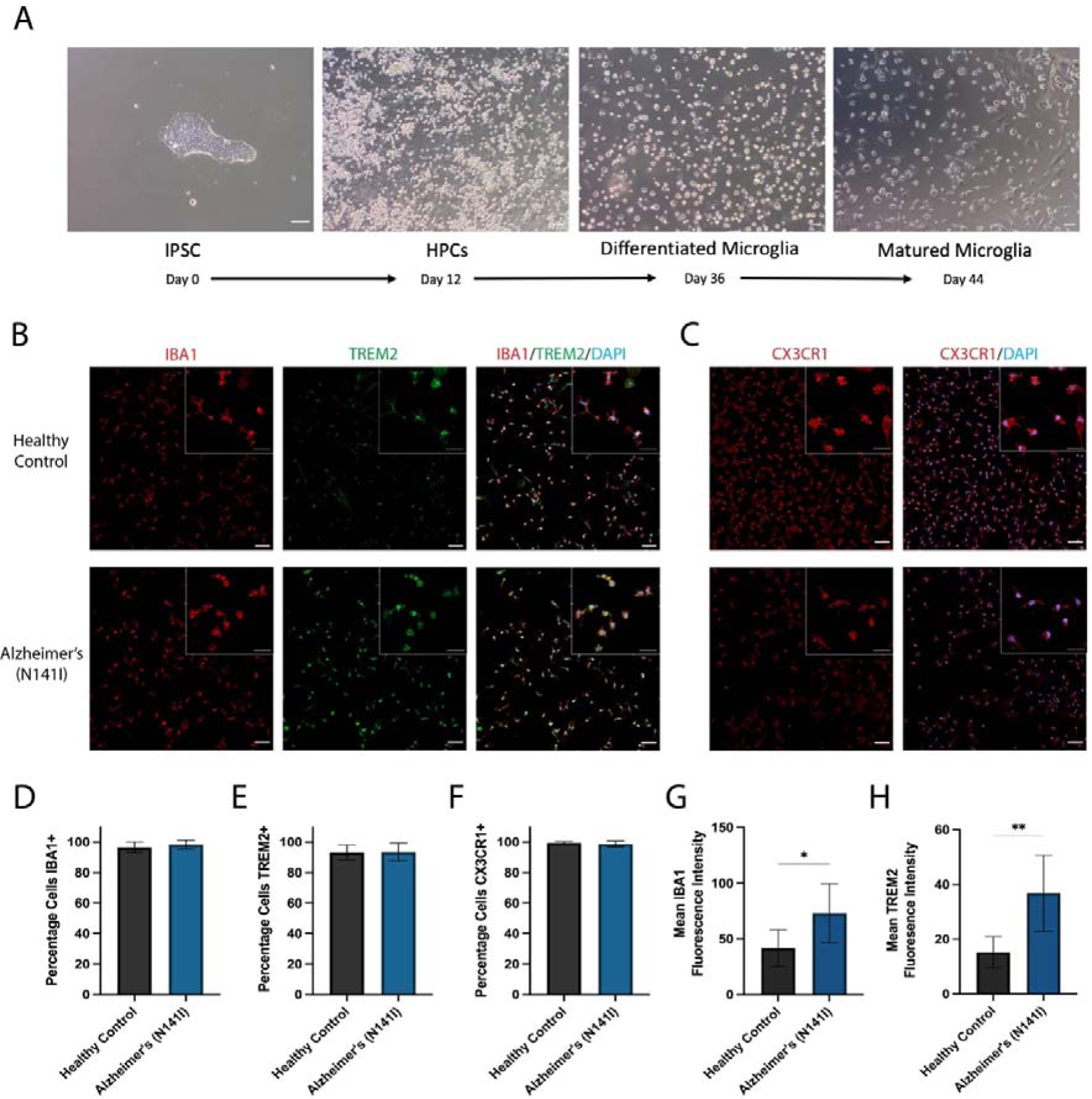
A) Overview of differentiation timeline for iPSC-derived microglia-like cells showing representative brightfield images on the last day of each stage. B) Representative immunofluorescence images of iPSC-derived astrocytes from healthy control lines and familial AD lines harbouring a PSEN2 (N141I) mutation. The cells were stained for microglial markers IBA1 (red), TREM2 (green) and C) CX3CR1 (red) and all nuclei were counterstained with DAPI (blue). Insert shows higher magnification. All scale bars = 50 μm. The percentage of iPSC-derived microglia-like cells positive for D) IBA1, E) TREM and F) CX3CR1. Mean fluorescence intensity of G) IBA1 and H) TREM2 per cell. An unpaired t-test was used to test whether there were statistically significant differences between the means of AD-derived and healthy control astrocytes. Each bar displays the mean ± SD of three cell lines with n ≥ 2 independent experiments per line. (** p < 0.01). Immunofluorescence staining of all cell lines can be found in Additional file 1: Fig. S4.

The results presented herein reveal that the fAD-causative PSEN2 (N141I) mutation did not affect the ability of iPSCs to successfully differentiate into mature astrocytes or microglia-like cells *in vitro*. Although, the expression of the astrocytic proteins GFAP and S100β were shown to be upregulated in PSEN2 (N141I)-mutant astrocytes.

### AD-Derived Astrocyte Morphology is Atrophied, Less Heterogenous and More Activated

Previous reports of iPSC-derived astrocytes from both sporadic and fAD patients have shown these cells exhibit substantial changes to the morphological phenotype compared to astrocytes derived from healthy control iPSCs (23). Hence, we quantified the morphological profiles of iPSC-derived astrocytes from our cohort of fAD patients and healthy controls. AD-derived PSEN2 (N141I)-mutant astrocytes displayed a significant reduction in the average perimeter of each cell compared to healthy controls (142.1 μm^2^ ± 47.5 SD and 409.1 μm^2^ ± 212.7 SD, respectively) (p < 0.05; Fig. 4 A). Cellular circularity is a quantitative index reflecting the extent of process extension, with a measurement of 1 describing a perfectly round cell. AD-derived astrocytes exhibited a rounder, more spherical shape in comparison to healthy control astrocytes, as shown by a significant increase in cellular circularity (0.55 ± 0.09 SD and 0.31 ± 0.10 SD, respectively) (p < 0.001; Fig. 4 B). Three-dimensional reconstruction and determination of cell volume was conducted using fine z-stacks, finding AD-derived astrocytes to have a significant reduction in average cell volume compared to healthy controls (7653 μm^3^ ± 2889 SD and 21126 μm^3^ ± 9911 SD, respectively) (p < 0.01; Fig. 4 C).

**Fig. 4.**
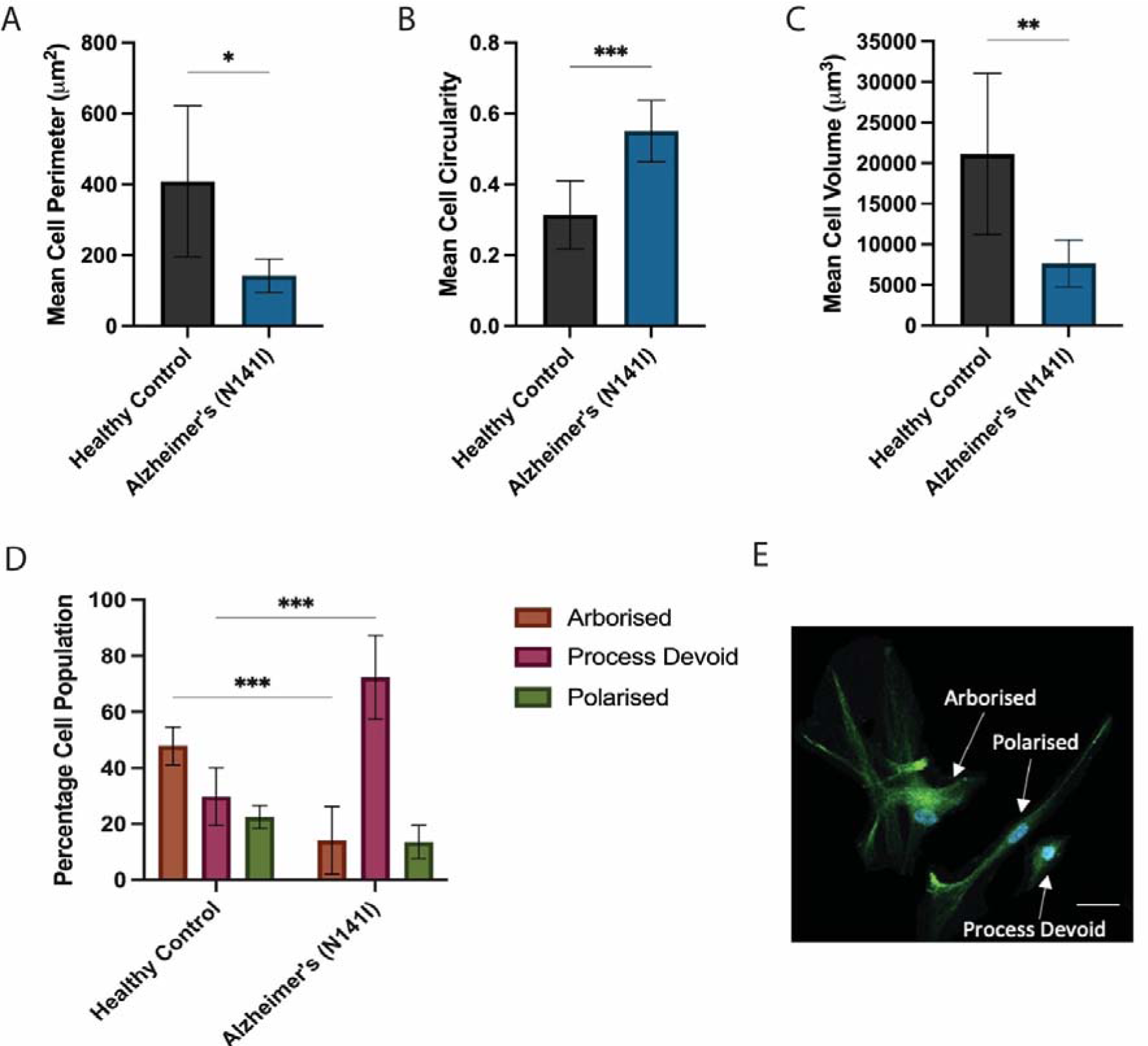
Morphological characterisation of iPSC-derived astrocytes from three healthy control and three familial AD lines harbouring a PSEN2 (N141I) mutation. Quantification of A) average cell perimeter, B) average cell circularity and C) average cell volume were determined using GFAP and S100β immunofluorescence and 3D reconstruction within FIJI. (C) Individual cells were categorised and based on morphological appearance, with representative phenotypes of ‘arborised’, ‘polarised’ and ‘process devoid’ cells illustrated in E). An unpaired t-test (A-C) or two-way ANOVA with a post-hoc Dunnett’s test (D) were used to test whether there were statistically significant differences between the means of AD-derived and healthy control astrocytes for each morphological category. The figure displays the mean ± SD of three cell lines with n ≥ 2 independent experiments per line (* p < 0.05, ** p < 0.01, *** p < 0.001). Scale bar = 50 μm.

To examine how morphological distribution of astrocytes varied across the cellular population, we classified individual astrocytes into various morphological categories. These categories included ‘arborised’, ‘polarised’ and ‘process devoid’ astrocyte morphologies, previously defined by Jones *et al.,* (23). Healthy control astrocytes most commonly presented as large cells with multiple lengthy, branching processes which is highly characteristic of an archetypal astrocyte morphology, referred to as ‘arborised’ (47.9 % ± 6.7 SD) (Fig. 4 D). The remaining cells were similarly split between bipolar-type cells protruding two main processes, referred to as ‘polarised’ (22.5 % ± 4.1 SD), and smaller cells with small to no extensions protruding from the cell body, referred to as ‘process devoid’ (29.7 % ± 10.3 SD) (Fig. 4 D). Representative images of each morphological classification are shown in Fig. 4 E. In contrast, AD-derived astrocytes showed a general reduction in cellular heterogeneity compared to healthy controls. Specifically, AD astrocytes were predominantly comprised of process-devoid cells (72.3 % ± 14.9 SD), which constituted a significantly higher proportion of cells compared to healthy controls (p < 0.001; Fig. 4 D). The proportion of AD-derived astrocytes that exhibited the archetypal arborised morphology was significantly lower than healthy controls (14.2 % ± 12.1 SD) (p < 0.001; Fig. 4 D), however there was no significant difference between the proportion of polarised cells (Fig. 4 D).

### AD-Derived Microglia Show Less Complex Ramification

Reduction in the extent and complexity of microglial ramification isolated from the post-mortem brain tissue of AD patients has been previously reported (36, 37). These changes in morphological structure are hypothesised to parallel changes in AD-associated microglial functions. Hence, we performed a quantitative analysis of iPSC-derived microglial-like cell morphology and characterised cellular ramifications through the number, length and complexity of branching processes. The automated analysis of immunofluorescence images presented herein closely followed a previously developed protocol (38). A representative image post-processing and skeletonization is shown in Fig. 5 A. AD-derived microglia-like cells showed a significant reduction in the average number of extension branch points, termed junctions, compared to healthy controls (5.9 ± 1.2 SD and 13.4 ± 7.8 SD, respectively) (p < 0.05, Fig. 5 B). Similarly, AD-derived microglia-like cells showed a significant decrease in the average number of branches compared to healthy controls (12.5 ± 2.6 SD and 25.9 ± 13.8 SD, respectively) (p < 0.05, Fig. 5 C). Another key metric of ramification includes the number of contact points at the end of each branch, termed endpoints. We found AD-derived microglia-like cells to have significantly fewer endpoints per cell compared to healthy controls (6.8 ± 1.4 and 11.0 ± 3.6 SD, respectively) (p < 0.05, Fig. 5 D). To gauge how far microglial processes extended from the cell body we analysed the average longest branch length per cell and found no significant difference between AD-derived microglia-like cells and healthy controls (11.6 μm ± 1.9 SD and 13.3 μm ± 4.7 SD, respectively) (Fig. 5 E). Taken together, these results demonstrate that PSEN2 (N141I)-mutant astrocytes and microglia-like cells show less complex ramified morphology.

**Fig. 5.**
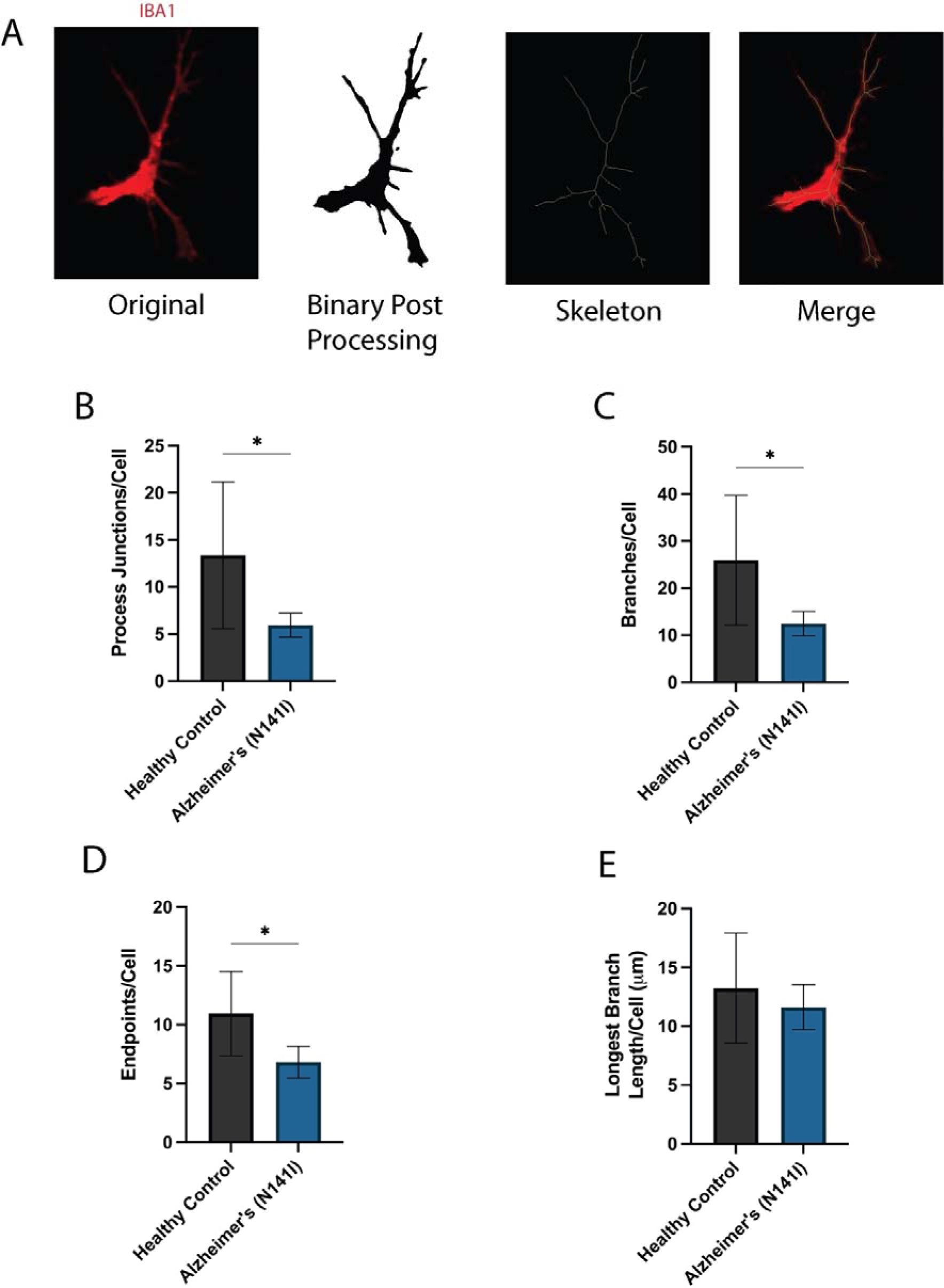
A) An example of the processing of an immunofluorescence image of iPSC-derived microglia-like cells stained for IBA1 (red), the result of image processing is shown in binary and was followed by skeletonization (green). An overlay of the original IBA1 image (red) and skeleton (green) is shown for comparison. Automated analysis of the mean number of B) junctions, C) branches, D) endpoints and E) longest branch length per cell. An unpaired t-test was used to test whether there were statistically significant differences between the means of AD-derived and healthy control microglia-like cells. The figure displays the mean ± SD of three cell lines with n = 2 independent experiments per line (* p < 0.05).

### AD-Derived Astrocytes and Microglia Exhibit an Exaggerated Pro-Inflammatory Response to Immune Challenge

We next aimed to functionally characterise these microglia-like cells and astrocytes in terms of cytokine and chemokine secretion. Firstly, we wanted to confirm that our iPSC-derived astrocytes would respond to LPS stimulation and release IL-6, a signature that we would expect to see from mature astrocytes. Both the healthy control and AD-derived astrocytes basally secreted small amounts of IL-6 (26.9 pg/mL ± 29.4 SD and 74.1 pg/mL ± 83.5 SD, respectively) (Fig. 6 A) and showed increased secretion of IL-6 from basal upon stimulation with 10 µg/mL LPS (454.5 pg/mL ± 589.3 SD and 398.3 pg/mL ± 387.7 SD, respectively) and 50 µg/mL LPS (460.6 pg/mL ± 552.3 SD and 478.0 pg/mL ± 447.4 SD, respectively), albeit no significant differences being found between healthy controls and AD-derived astrocytes (Fig. 6 A).

**Fig. 6.**
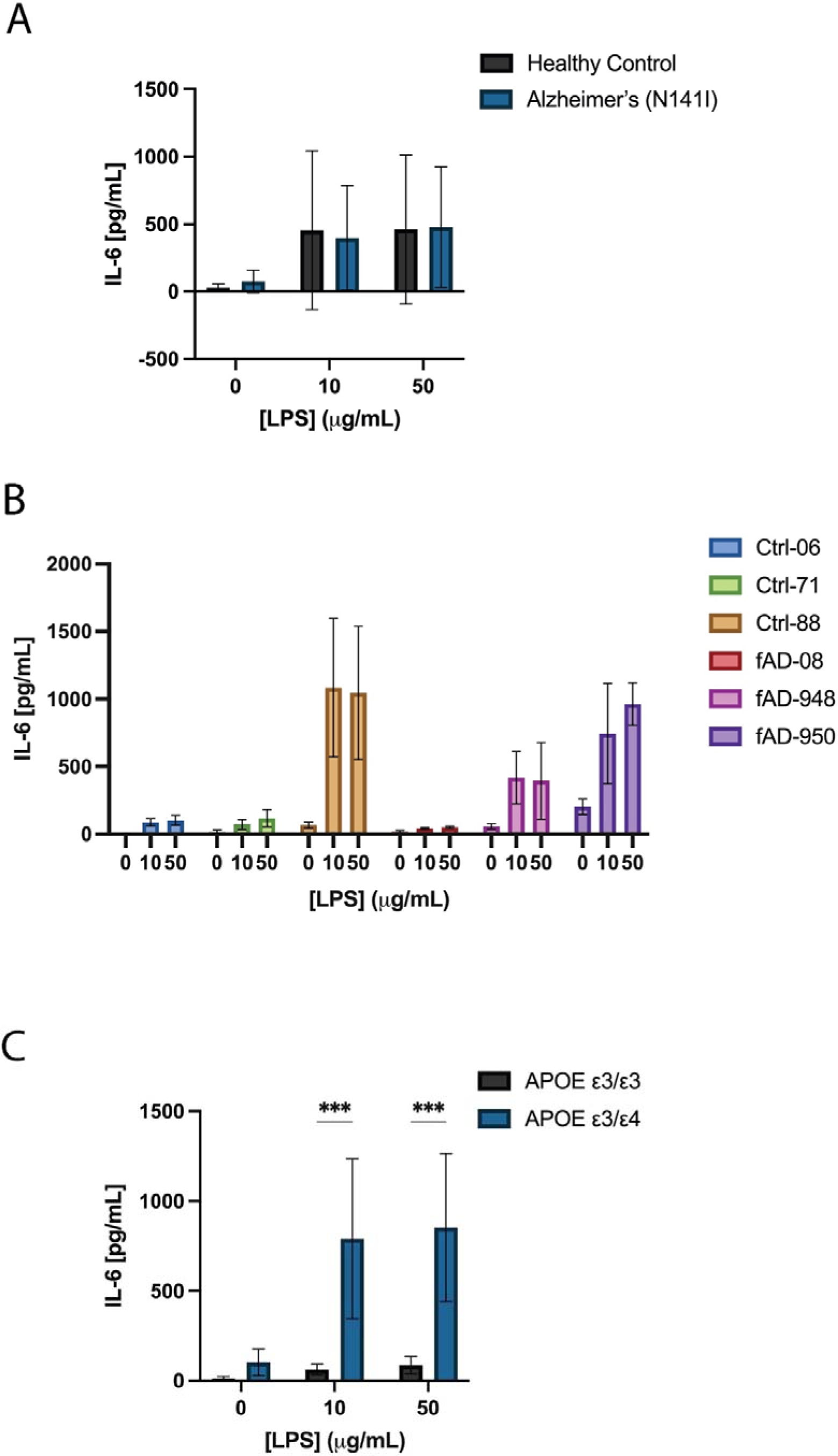
Concentration of IL-6 secreted from healthy control or AD-derived astrocytes A) basally and after exposure to 10 or 50 μg/mL LPS for 24 h. B) Data from part A represented as individual cell lines. Cell lines were re-categorised based on their APOE ε3/ε3 (Ctrl-06, Ctrl-71 and fAD-08) or APOE ε3/ε4 (Ctrl-88, fAD-948 and fAD-950) genotype and shows the concentration of IL-6 secretion C) basally and after exposure to 10 or 50 μg/mL LPS for 24 h. A two-way ANOVA with a multiple comparisons and post-hoc Dunnett’s test was used to test whether there were statistically significant differences between the means of AD-derived and healthy control astrocytes or APOE ε3/ε3 and APOE ε3/ε4 astrocytes (*** p < 0.001). Statistical tests were not conducted for figure B. Figures A and C display the mean ± SD of three cell lines with n ≥ 2 independent experiments per line, B display the mean ± SD of n ≥ 2 independent experiments per line.

A large amount of variability was observed in both the healthy control and AD-derived groups in the basal and LPS-stimulated conditions (Fig. 6 A). In an attempt to identify a source of the large variation in IL-6 secretion, we reorganised the grouped results into their individual cell lines. The Ctrl-88, fAD-948 and fAD-950 cell lines secreted higher IL-6 basally and showed, on average, a 9.2-fold higher response to 10 µg/mL LPS and 7.2-fold to 50 µg/mL LPS compared to the Ctrl-06, Ctrl-71 and fAD-08 lines (Fig. 6 B). Further investigation of the genotypes of these three high responding lines (Ctrl-88, fAD-948 and fAD-950) revealed they were heterozygous carriers of the APOE ε4 allele, the largest associated risk gene for sAD. Conversely, the three lower responders (Ctrl-06, Ctrl-71 and fAD-08) were homozygous for the APOE ε3 allele (Table 1). This led us to perform a post-hoc analysis where we recategorized cell lines into their respective APOE genotype, being ε3/ε3 or ε3/ε4. We found a non-significant increase in basal IL-6 (ε3/ε3: 13.8 pg/mL ± 9.9 SD and ε3/ε4: 101.4 pg/mL ± 73.7 SD) (Fig. 6 C) and significant increases in 10 µg/mL LPS-stimulated (ε3/ε3: 63.6 pg/mL ± 29.4 SD and ε3/ε4: 789.2 ± 444.6 SD) (p < 0.001; Fig. 6 C) and 50 µg/mL LPS-stimulated IL-6 secretion (ε3/ε3: 87.1 pg/mL ± 48.9 SD and ε3/ε4: 851.5 ± 411.1 SD) (p < 0.001; Fig. 6 C) from APOE ε3/ε4 astrocytes compared to APOE ε3/ε3. It is important to note that the statistical test here is not a *bona fide* reflection of statistical significance due to the post-hoc nature of the analysis. Nevertheless, it does provide a hypothesis to explain the variability originally observed when comparing IL-6 secretion from healthy control and AD-derived astrocytes. Furthermore, it provided an important consideration when interpreting results from subsequent IL-6 ELISAs.

In addition to exhibiting neurotoxic properties, Aβ_42_ is known to elicit glial activation and causes release of pro-inflammatory cytokines (39). As such, we aimed to test the effect that an AD-relevant stimulus has on our iPSC-derived astrocytes and microglia-like cells. We first characterised the IL-6 response from iPSC-derived astrocytes after being exposed to both 5 and 10 μM monomeric Aβ_42_. AD-derived astrocytes showed a significantly elevated secretion of IL-6 compared to healthy controls after exposure to both 5 μM (963.2 ± 697.6 SD and 130.0 ± 85.9 SD, respectively) (p < 0.01) and 10 μM Aβ_42_ (1023.0 pg/mL ± 621.7 SD and 118.8 pg/mL ± 90.9 SD, respectively) (p < 0.001; Fig. 7 A). As Aβ_42_ has been shown to be neurotoxic in *in vitro* cultures of both primary and iPSC-derived neurons (40, 41) we aimed to test whether this toxicity extended to iPSC-derived astrocytes. Neither healthy control or AD-derived astrocytes showed any reduction in cell viability from basal after 5 μM (110.9 % ± 15.7 SD and 104.2 % ± 3.9 SD, respectively) or 10 μM Aβ_42_ exposure (108.0 % ± 24.0 SD and 107.3 % ± 18.9 SD, respectively) (Fig. 7 B). This provided confidence that the secreted IL-6 was a cellular response and not an artefactual release from apoptotic cells. In light of the previous results found with LPS-stimulated IL-6 secretion and APOE genotype, we conducted a similar post-hoc analysis recategorizing the cells into their respective APOE ε3/ε3 and APOE ε3/ε4 genotypes. In contrast to LPS stimulation, we observed a larger amount of variation after reorganisation and no significant differences between the APOE genotypes in IL-6 secretion (Fig. 7 C) or cell viability (Fig. 7 D) were found after stimulation with 5 or 10 μM Aβ_42._

**Fig. 7.**
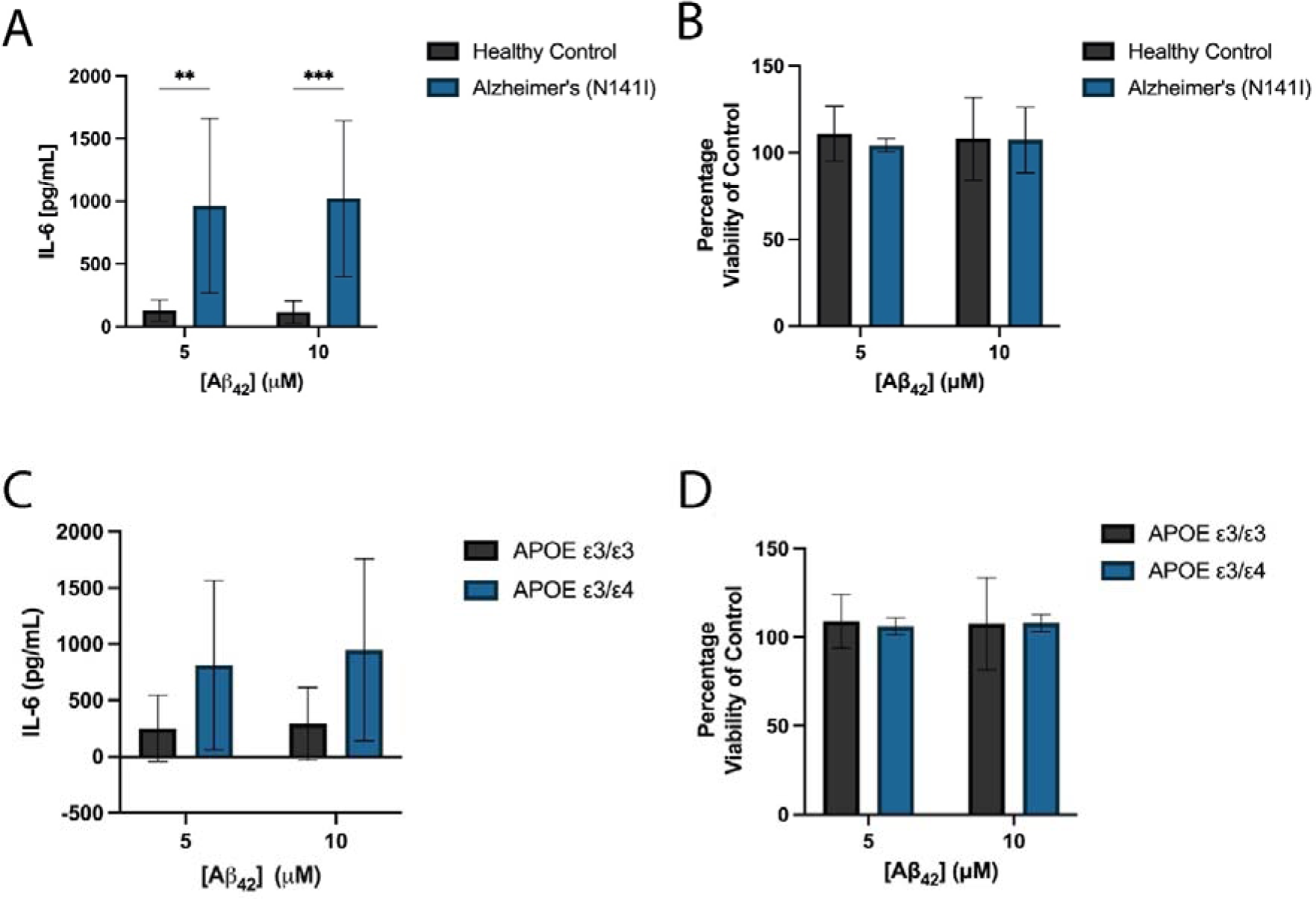
The concentration of A) secreted IL-6 and B) viability of AD-derived and healthy control astrocytes after stimulation with initially monomeric 5 or 10 μM Aβ_42_ for 24 h. Cell lines were re-categorised based on their APOE ε3/ε3 (Ctrl-06, Ctrl-71 and fAD-08) or APOE ε3/ε4 (Ctrl-88, fAD-948 and fAD-950) genotype and shows the concentration of IL-6 secretion E) basally and F) after exposure to 5 or 10 μM Aβ_42_ for 24 h. The figure displays the mean ± SD of three cell lines with n ≥ 2 independent experiments per line. A two-way ANOVA with post-hoc Dunnett’s test was used to test whether there were statistically significant differences between mean of AD-derived and healthy control astrocytes and APOE ε3/ε3 and APOE ε3/ε4 astrocytes (** p < 0.01, *** p < 0.001).

After observing significant differences in the release of IL-6 between AD-derived and healthy control-derived astrocytes, we conducted a broader characterisation of cytokines/chemokines from iPSC-derived astrocytes and microglia-like cells using a multi-cytokine array consisting of 36 cytokines/chemokines (Additional File 1: Fig. S5). We observed no significant changes in the basal secretion of cytokines between healthy and AD-derived astrocytes (Fig. 8 A). Upon stimulation with 10 μM Aβ_42_ we observed significant increases in the secretion of chemokine (C-X-C motif) ligand 1 (CXCL1) (2974 a.u ± 2677 SD and 10772 a.u ± 1156 SD, healthy control and AD-derived respectively) (p < 0.01), intercellular adhesion molecule 1 (ICAM-1) (871 a.u ± 185 SD and 4533 a.u ± 1675 SD, healthy control and AD-derived respectively) (p < 0.01), IL-6 (968 a.u ± 426 SD and 5297 a.u ± 4814 SD, healthy control and AD-derived respectively) (p < 0.01) and IL-8 (5297 a.u ± 4814 SD and 21540 a.u ± 4946 SD, healthy control and AD-derived respectively) (p < 0.01) from the AD-derived astrocytes compared to healthy controls (Fig. 8 B). The significant increase in IL-6 mirrors that of the specific IL-6 ELISA previously presented (Fig. 7 A) and provides confidence that this lower sensitive multi-cytokine array was detecting differences in moderately low cytokine concentrations.

**Fig. 8.**
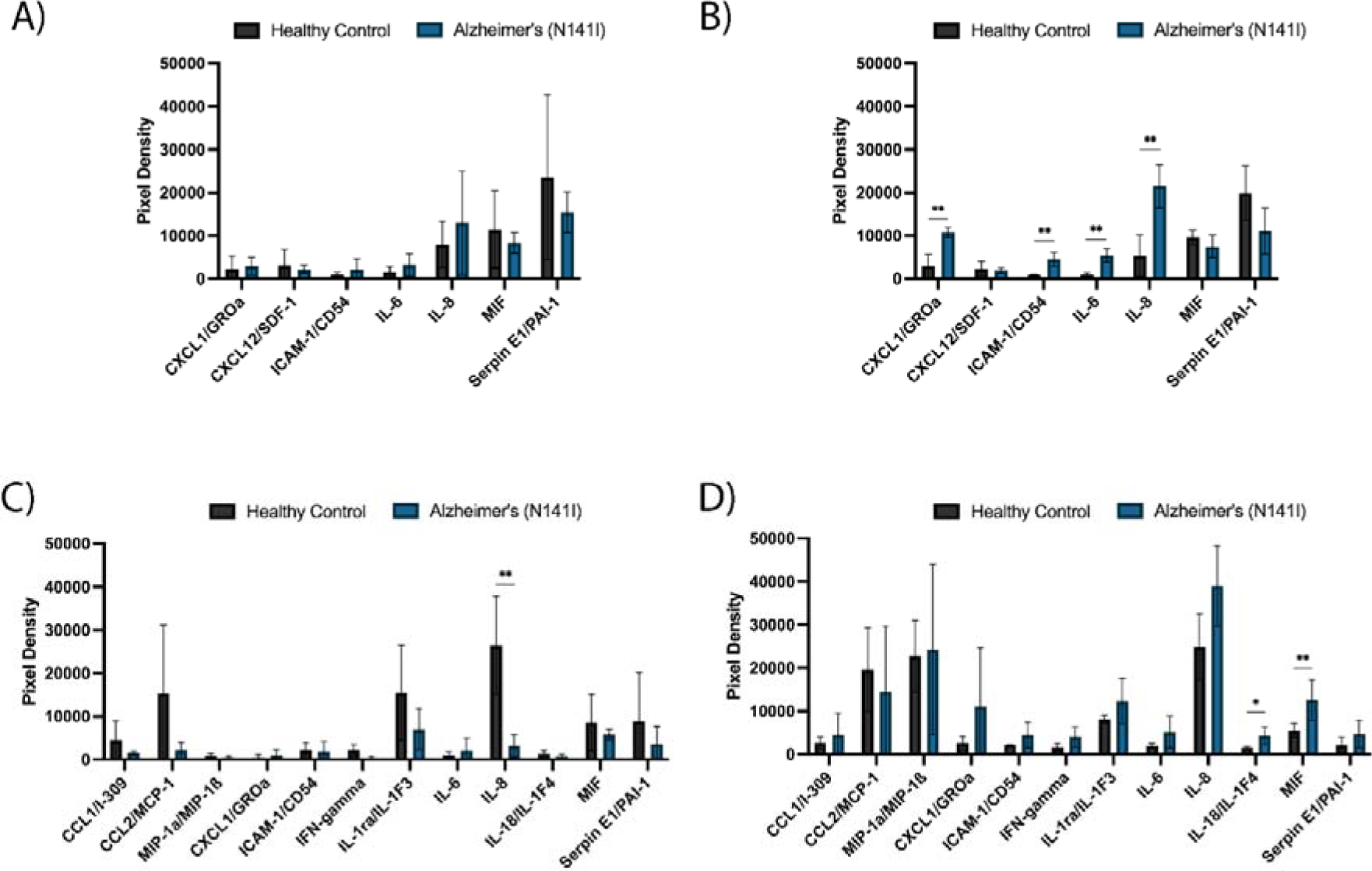
Multi-cytokine array of AD-derived or healthy iPSC-derived astrocytes A) basally and B) after 24 h exposure to 10 μM Aβ_42_ and iPSC-derived microglia-like cells C) basally and D) after 24 h exposure to 10 μM Aβ_42_. The figure displays the mean ± SD of three cell lines with the average of two experimental duplicates per line. Multiple unpaired, non-parametric Mann-Whitney t-tests adjusting for a 5 % false discovery rate were used to test whether there were statistically significant differences between mean cytokine/chemokine release of AD-derived and healthy control astrocytes and microglia-like cells (* p < 0.05, ** p < 0.01). Cytokines/chemokines with mean pixel density under 10 % of the maximum in both basal and Aβ_42_-stimulated conditions were considered background and are not shown, full cytokine array is shown in Additional file 1: Fig. S5.

iPSC-derived microglia-like cells from AD origin showed a significant reduction in the basal secretion of IL-8 compared to healthy controls (26457 a.u ± 11408 SD and 3148 a.u ± 2582 SD, respectively) (p < 0.01; Fig. 8 C). Upon stimulation with Aβ_42_, AD-derived microglia-like cells showed a considerable increase in IL-8 (12.4-fold from basal), whereas the IL-8 secreted from healthy control microglia-like cells stayed relatively unchanged (1.1-fold decrease) (Fig. 8 D). Significantly increased secretion of IL-18 (1473 a.u ± 352 SD and 4299 a.u ± 2006 SD, healthy control and AD-derived respectively) (p < 0.05) and macrophage migration inhibitory factor (MIF) (5483 a.u ± 1635 SD and 12535 a.u ± 4742 SD, healthy control and AD-derived respectively) (p < 0.01) were observed from AD-derived microglia-like cells compared to healthy controls after Aβ_42_ stimulation (Fig. 8 D). We observed a substantial increase in macrophage inflammatory protein-1 (MIP-1) secretion after Aβ_42_ exposure for both the healthy control (27.1-fold increase from basal) and AD-derived microglia-like cells (44.5-fold increase from basal), however there were no significant differences between the final levels of secretion between AD-derived and healthy control microglia-like cells.

In addition to the release of cytokine/chemokines, astrocytes not only secrete Aβ, but are suggested to do so in a high enough quantity to contribute to Aβ load within the brain (42). Despite this, the effect of fAD-causative mutations on Aβ_42_ production has been primarily studied in neurons. Therefore, we analysed the secretion of the AD-associated peptides Aβ_42_ and Aβ_40_ 72 hours post-plating of our control and AD-derived astrocytes harbouring a PSEN2 (N141I) mutation. AD-derived astrocytes produced a significant increase in the secretion of Aβ_42_ compared to healthy controls (0.0600 pg/μg ± 0.0129 SD and 0.0372 pg/μg ± 0.0207 SD, respectively) (p < 0.05; Fig. 9 A). No significant differences were observed in the secretion of Aβ_40_ (0.317 pg/μg ± 0.076 SD and 0.276 pg/μg ± 0.111 SD, AD-derived and healthy control respectively) (Fig. 9 B), the ratio of Aβ_42:40_ (0.177 ± 0.068 SD and 0.127 ± 0.023 SD, AD-derived and healthy control respectively) (Fig. 9 C) or total level of Aβ secreted (0.362 pg/μg ± 0.079 SD and 0.322 pg/μg ± 0.144 SD, AD-derived and healthy control respectively) (Fig. 9 D) between AD-derived astrocytes and healthy controls. Additionally, we found no differences in the levels of Aβ_42_, Aβ_40_, the Aβ_42:40_ ratio and total Aβ between APOE ε3/ε3 and APOE ε3/ε4 astrocytes (Additional file 1: Fig. S6).

**Fig. 9.**
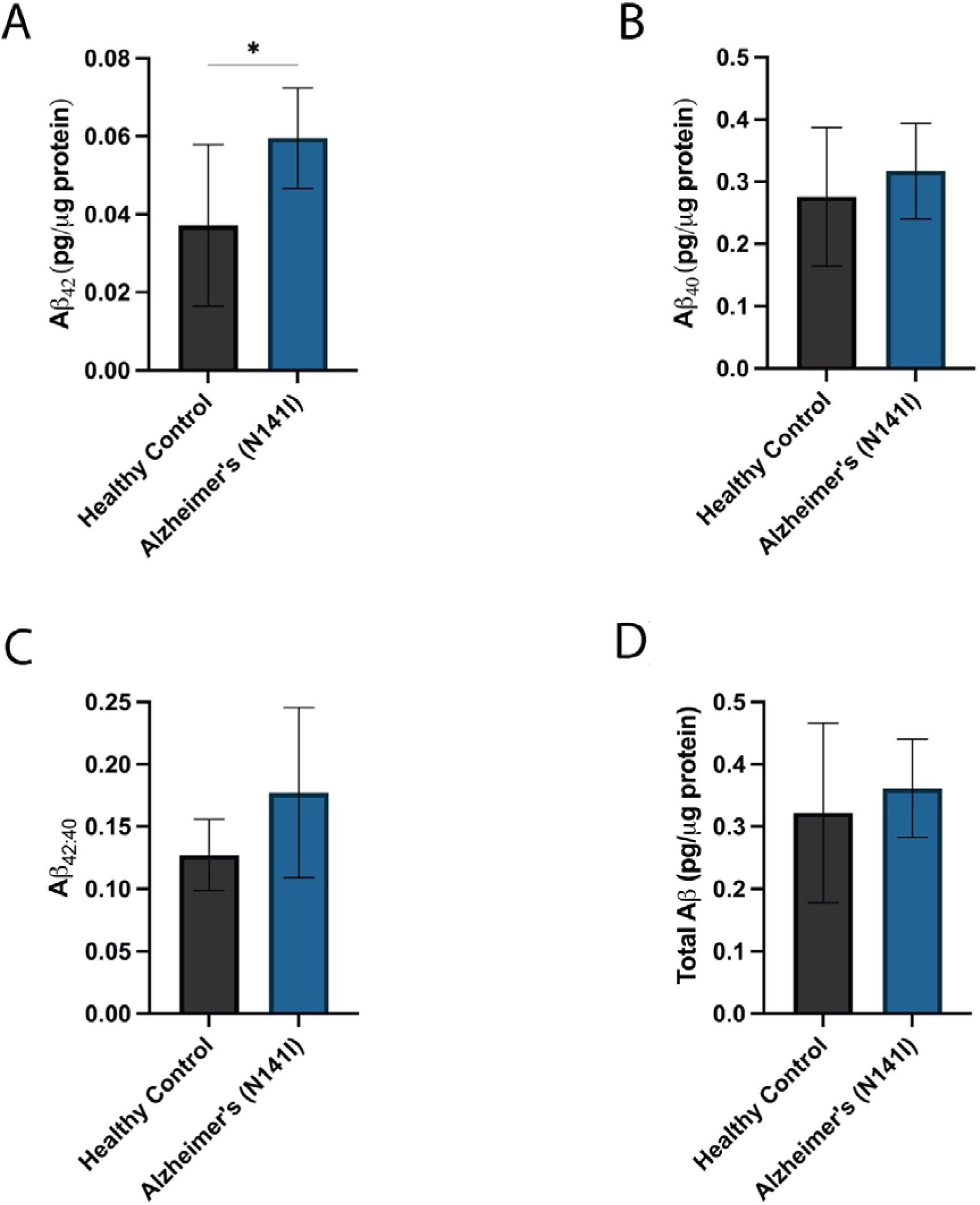
Concentrations of A) Aβ_42_, B) Aβ_40_, C) the ratio of Aβ_42:40_ and D) total Aβ protein quantified from iPSC-derived astrocyte supernatants 72 h after plating. Secreted Aβ concentrations were measured using a highly-sensitive ELISA and normalised to total protein concentration determined by BCA. The figure displays the mean ± SD of three cell lines with n ≥ 2 independent experiments per line. An unpaired t-test was used to test whether there were statistically significant differences between mean of AD-derived and healthy control astrocytes (* p < 0.05).

### Astrocyte and Microglial Phagocytosis of Aβ_42_

Phagocytosis is one of the major functions of CNS glial cells. To assess the phagocytic capacity of AD-derived and healthy control astrocytes and microglia-like cells, we exposed the glial cells to fluorescently tagged fibrillar Aβ_42_ based on a previously published protocol using iPSC-derived microglia (43). AD-derived astrocytes on average exhibited significantly greater uptake of fluorescent Aβ_42_ fibrils per cell compared to healthy controls (1.33 × 10^7^ ± 8.67 × 10^6^ SD and 3.84 × 10^6^ ± 3.62 × 10^6^ SD, respectively) (p < 0.05; Fig. 10 A). The proportion of astrocytes that showed Aβ_42_ uptake was very high (> 95 %) in both the AD-derived and healthy control lines, finding no statistical differences between each group (98.7 % ± 1.5 SD and 97.4 % ± 3.3 SD, respectively) (Fig. 10 B). Similarly, iPSC-derived microglia-like cells showed a very high proportion of phagocytic cells (> 95 %) with no significant differences between AD-derived and healthy controls (97.5 % ± 2.2 SD and 98.5 % ± 0.9 SD, respectively) (Fig. 10 D). Surprisingly, we observed no significant differences between the amount of Aβ_42_ taken up by AD-derived or healthy control microglia-like cells (2.71 × 10^6^ ± 1.30 × 10^6^ SD and 1.64 × 10^6^ ± 1.15 × 10^6^ SD, respectively) (Fig. 10 C). Representative images of iPSC-derived astrocytes (Fig. 10 E) and microglia-like cells (Fig. 10 F) showcases the strong uptake of fluorescent Aβ_42_ and high proportion of phagocytic cells.

**Fig. 10.**
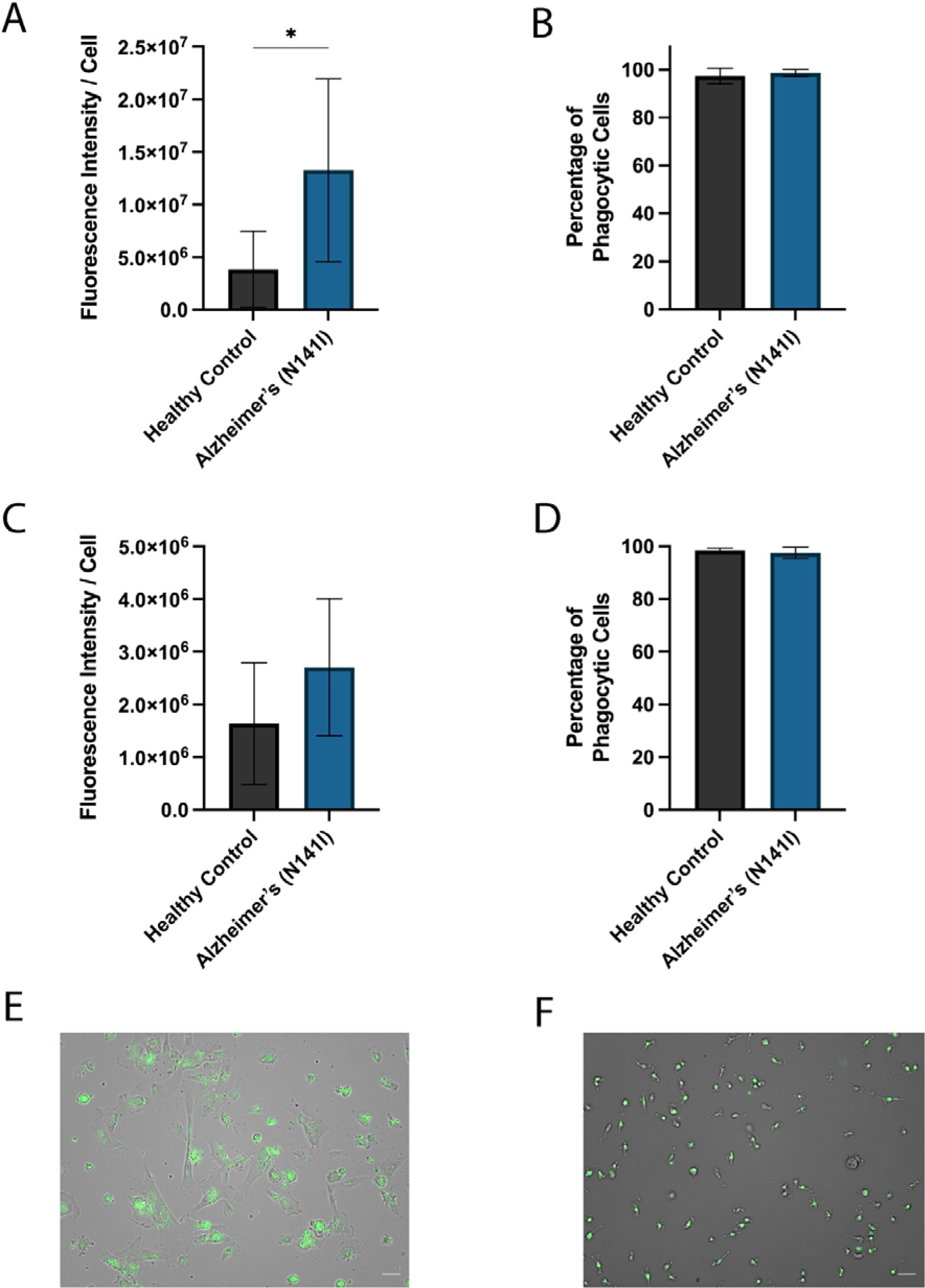
AD-derived and healthy iPSC-derived astrocytes were exposed to 488-fluorescent-Aβ_42_ fibrils for 2 h and phagocytosis was quantified through average fluorescent intensity per cell for A) iPSC-derived astrocytes and C) iPSC-derived microglia-like cells. The figure shows the percentage of B) iPSC-derived astrocytes and D) iPSC-derived microglia-like cells which show any uptake of Aβ_42_. The figure displays the mean ± SD of three cell lines with n ≥ 2 independent experiments per line. An unpaired t-test was used to test whether there were statistically significant differences between mean of AD-derived and healthy control astrocytes (* p < 0.05). Representative overlay of brightfield and fluorescent images of E) iPSC-derived astrocytes and F) iPSC-derived microglia-like cells, fluorescent Aβ_42_ fibrils shown in green, scale bar = 50 μm.

## Discussion

Astrocytes and microglia play a crucial role in many aspects of AD pathophysiology such as neuroinflammation and dysregulation of Aβ proteostasis. However, the specific cellular dysfunction occurring and the molecular drivers of glial activation during various stages of AD progression is not well understood. Here we describe the previously unreported differentiation and characterisation of iPSC-derived astrocytes and microglia-like cells from fAD patients harbouring a PSEN2 (N141I) mutation. We showed that PSEN2 (N141I)-mutant astrocytes and microglia-like cells presented with an extensive disease-associated phenotype with reduced morphological complexity, exaggerated pro-inflammatory cytokine secretion and altered Aβ_42_ production and phagocytosis (Summarised in Fig. 11).

**Fig. 11.**
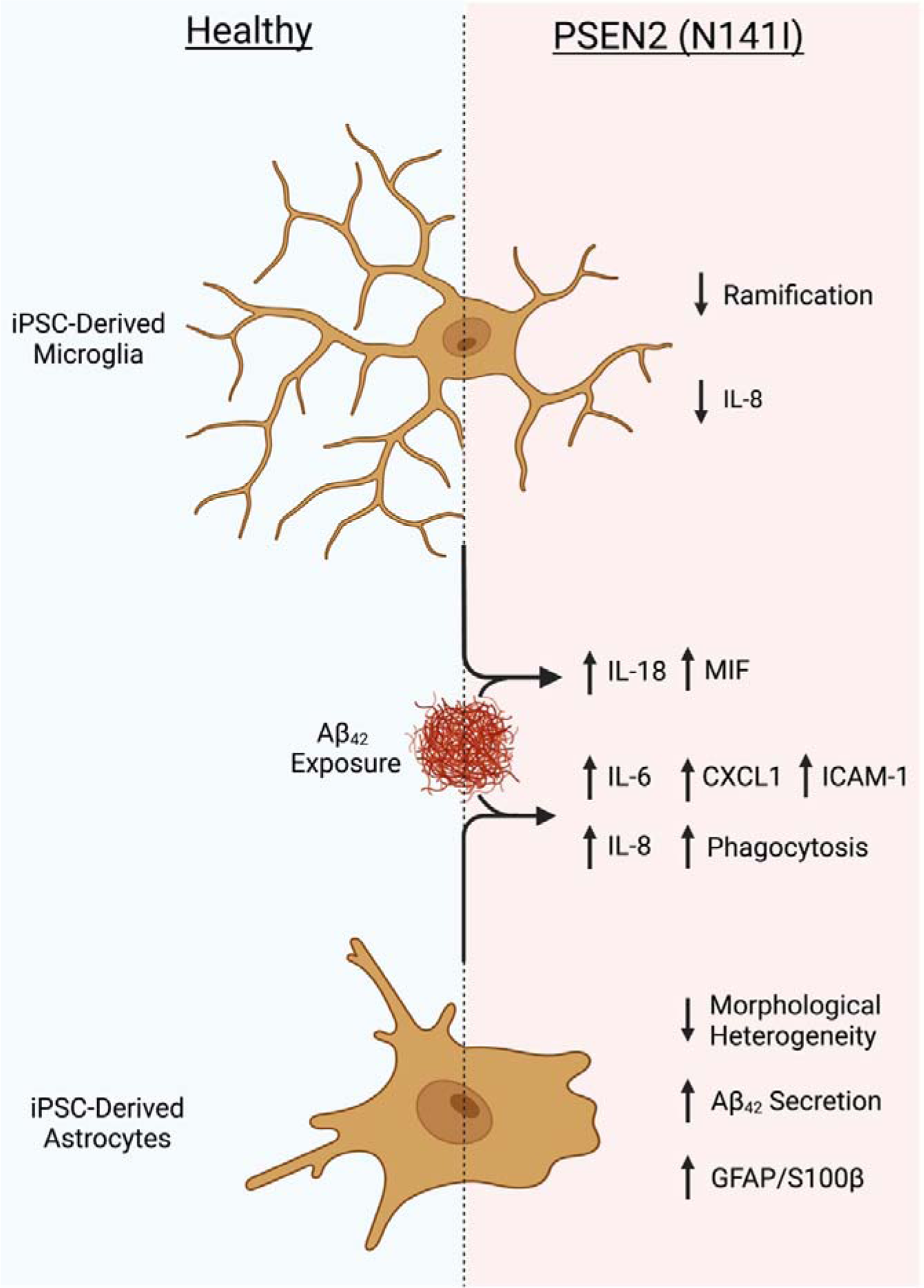
Graphical representation and summary of morphological and immunological features of iPSC-derived microglia-like cells (top) and astrocytes (bottom) from healthy (left) and PSEN2 (N141I)-mutant fAD (right) origin. Created with Biorender.com

It is worthy of note that classifying *in vitro* iPSC-derived microglia-like cells as *bona fide* microglia is a somewhat contentious topic within literature and is generally supported with extensive transcriptomic analysis (21). Despite our differentiation following a previously characterised method and producing cells positive for markers strongly consistent with microglial identity (IBA1, TREM2 and CX3CR1) (44), we cannot conclusively infer *bona fide* microglial identity based solely on immunofluorescence staining as these proteins can be expressed on other myeloid cells such as monocytes and macrophages, hence our reference to these cells as microglia-like cells.

Changes in microglial and astrocyte morphology have been observed in the brain of AD patients and can reveal aspects about cellular phenotypes and activation states (45, 46). Cell autonomous morphological changes in our AD-derived astrocytes mirror findings from iPSC-derived astrocytes from both sAD and fAD PSEN1-mutant origin, finding similar changes to astrocyte size, shape and morphological distribution (23). Furthermore, triple transgenic and APP-mutant mouse models of AD also show reduction to astrocyte surface area, volume and process complexity, often before the onset of Aβ accumulation (47-49). This suggests that the morphological changes associated with AD-derived astrocytes observed within the current study are not specific to the PSEN2 (N141I) mutation, but rather a common cell-autonomous dysfunction across multiple familial and sporadic origins which mirror those of activated pro-inflammatory iPSC-derived astrocytes (50). In addition to morphological changes, GFAP is widely used as a marker for astrocyte activation (51) and is found to be upregulated in the brains of AD patients along with S100β (52-57). Alongside the morphological characteristics, the increase in GFAP and S100β expression found within our AD-derived astrocytes suggests a basal skew towards a more activated phenotype.

Similar to AD-derived astrocytes, iPSC-derived microglia-like cells from fAD patients showed a reduction in the number and complexity of processes extending from the glial cell body, however the length in which the processes are extending from the cell body seemed to be unchanged between disease groups. Various microglial morphologies have been defined using human post-mortem brain tissue, with each attributed to varying levels of activation (58). It is suggested that under physiological conditions, microglia present as ramified cells with highly branched processes extending substantially from the cell body. Upon activation, microglia transition into a reactive morphology whereby the cells become less ramified through shorter and less complex processes (59), a characteristic observed in AD mouse models and in mice following ischemic injury (60-62). Similarly, PSEN2 (N141I) knock-in mice exhibited microglia with decreased ramification and number of dendritic branches, however no change in the branch length was observed (63, 64). Despite PSEN2 (N141I)-mutant microglia presenting with some morphological characteristics consistent with activation, namely reduced ramification, they do not entirely recapture the archetypal phenotype associated with microglial activation and suggest a more subtle change in phenotype. However, whether these morphological features are replicated in PSEN2 (N141I)-mutant AD patients is unknown as no morphometric analysis has been conducted on human brain tissue.

A large body of evidence has implicated neuroinflammatory changes as a key neuropathological feature within AD progression, largely associated with changes in the activation states of astrocytes and microglia (12, 65, 66). Unexpectedly, we found preliminary evidence that in iPSC-derived astrocytes the level of IL-6 secretion upon LPS stimulation was determined by APOE genotype and was independent of PSEN2 status. Within the CNS, APOE is a lipid transport protein acting as a ligand for low density lipoprotein receptor and primarily produced by astrocytes and is the highest genetic risk factor for sAD (1, 67). In humans, APOE presents as three allelic variants (APOE ε2, APOE ε3 and APOE ε4) and compared to individuals with the APOE ε3/ε3 genotype, the APOE ε3/ε4 genotype confers around a 3-fold increase in risk of developing AD and is significantly overrepresented in individuals with sAD (68, 69). A potential mechanism behind the disproportionate IL-6 response to LPS between APOE genotypes may lie within differences in the ability to bind to extracellular LPS. The single nucleotide polymorphism present within the APOE ε4 allele causes a reduction in the binding affinity of APOE to multiple extracellular proteins and LPS. As such, APOE ε3 binds to LPS with a higher affinity than the ε4 protein, reducing the ability of LPS to activate downstream pro-inflammatory cytokine production (70, 71). This hypothesis suggests that the presence of the ε4 allele does not directly increase the pro-inflammatory activity of LPS, but rather is less effective than the ε3 allele at negatively regulating LPS-mediated neuroinflammation. Astrocytes isolated from APOE ε4 mice exhibited reduced astrogliosis and IL-6 production following LPS stimulation compared to astrocytes isolated from APOE ε3 mice (72, 73). These findings are at odds with our data from iPSC-derived human astrocytes. This suggests that species differences may play a significant role in determining how the different isoforms of APOE mediate the response from astrocytes following neuroinflammatory stimulus.

We next examined the cytokine and chemokine profile of iPSC-derived astrocytes and microglia-like cells in response to exogenous Aβ_42_, a more AD-relevant stimulus than LPS. We found that fAD-derived astrocytes exhibited an exaggerated release of IL-6 in response to Aβ_42_, while variations in APOE genotypes had no significant effect. It is possible that any differences in the secretion of IL-6 between APOE genotypes within our cohort of iPSC-derived cells would be masked by the large effect observed with PSEN2 (N141I)-mutant lines. Future work utilising isogenic APOE lines will allow this to be investigated further.

Although IL-6 plays an important role in AD (74), the inflammatory profile of glial cells does not solely involve the level of IL-6 response, but rather the coordinated and unique secretion of multiple cytokines and chemokines. As such, the multi-cytokine array allowed us to analyse multiple cytokines simultaneously to gain a wider picture of the inflammatory profiles of AD-derived astrocytes and microglia-like cells. Both healthy and fAD-derived astrocytes basally secreted moderate levels of IL-8 while fAD-derived microglia-like cells exhibited a significant reduction in IL-8 secretion. Despite IL-8 being a pro-inflammatory cytokine (75), it also exhibits neurotrophic properties (76). Hence, reductions in microglial IL-8 secretion could contribute to AD progression and neuronal damage through a reduction in neural reparative mechanisms. Upon exposure to Aβ_42_, fAD-derived astrocytes exhibited significantly higher levels of ICAM-1, CXCL1, IL-6 and IL-8. In support of our findings, ICAM-1 is overexpressed by activated astrocytes in the brains of AD patients (77) and has been associated with Aβ pathology (78, 79). Blood brain barrier (BBB) dysfunctions have been identified as a key aspect of AD pathophysiology, to which astrocytes play an integral part (80). Previous literature has shown extracellular ICAM-1 alters tight-junction integrity and integrin adhesiveness in vascular endothelial cells, resulting in increased BBB permeability (81-83). A recent longitudinal study found elevated cerebrospinal fluid levels of ICAM-1, along with other cerebrovascular associated proteins, in the preclinical, prodromal and dementia stages of AD (84). Increased CXCL1 secretion was also found from Aβ-stimulated human astrocytes and was synaptotoxic *in vitro* (85). In addition to the BBB, astrocytes form an integral part of cortical synapses and the overproduction of CXCL1 may underlie astrocyte-mediated synaptotoxicity. Taken together with previous literature, our results provide evidence that astrocytes may contribute to cerebrovascular and synaptic dysfunction at an early stage within AD progression through altered ICAM-1 and CXCL1 secretion in response to extracellular Aβ.

Similar to astrocytes, fAD-derived microglia-like cells showed an exaggerated increase of IL-8 from basal in response to Aβ_42_. Significantly increased levels of the proinflammatory cytokines IL-18 and MIF upon Aβ_42_ exposure were unique to fAD-derived microglia-like cells. Our results are consistent with previous data reporting increased IL-18 secretion from activated mouse microglia (86) and increased levels of IL-18 and MIF within the brains and cerebrospinal fluid of AD patients, respectively (87, 88). In contrast to our findings, and these findings from human AD tissue, cultured microglia from PSEN2 (N141I) knock-in mice exhibited a different range of chemokine release in response to Aβ_42_ not impacted in our PSEN2 (N141I)-derived microglia such as the significant upregulation of CCL2, CCL5 and CXCL1 (63). Single-cell transcriptomic analysis highlight substantial species differences in microglial expression profiles and AD-associated genes such as PSEN (89, 90). As such, species differences may underlie the disparities between findings and emphasises the need to study human glial cells in the context of AD.

In addition to contributing to increased Aβ_42_ production, defective clearance of Aβ by glial cells is suggested to play a role in the progression of AD pathophysiology (91) and is supported by previous work finding reduced uptake of Aβ_42_ from PSEN1-mutant iPSC-derived astrocytes (26). However, we found fAD-derived astrocytes to uptake significantly more Aβ_42_ than healthy controls and suggest that phagocytic capacity may be impacted differentially by PSEN mutation status. The increased uptake of Aβ_42_ in our fAD-derived astrocytes may be another exaggerated immune response upon exposure to Aβ_42_, similar to our findings with Aβ_42_-stimulated cytokine release. How the change in Aβ_42_ uptake may contribute to AD progression is not clear. Perhaps the enhanced uptake of Aβ42 may be initially neuroprotective, however as the disease progresses the cells become dystrophic and unable to perform continuous immune functions. In contrast to other studies modeling AD using TREM2 knock-out iPSC-derived microglia (43, 92, 93), our results suggest that fAD-derived microglia do not exhibit cell-autonomous deficits in Aβ uptake. Additionally, the results presented herein mirror the phagocytosis of Aβ reported in APP and PSEN1-mutant iPSC-derived microglia (24) and other factors such as neuron-microglia or astrocyte-microglia communication are likely to impact microglial phagocytosis and could change over the course of disease progression.

Taken together, a common feature between fAD-derived astrocytes and microglia-like cells was the exaggerated release of multiple pro-inflammatory cytokines in response to Aβ_42_. Despite observing morphological characteristics in non-stimulated cells that were consistent with activated astrocytes and microglia, AD-derived glial cells exhibited elevated levels of pro-inflammatory cytokines only in response to Aβ_42_ and not basally. Our results suggests that unstimulated AD-derived astrocytes and microglia-like cells present with a primed phenotype, whereby the cells are predisposed to induce an exaggerated inflammatory response to a pathological stimulus.

## Conclusion

In conclusion, we report the novel and successful differentiation of astrocytes and microglia-like cells from iPSCs harbouring an fAD-causative PSEN2 (N141I) mutation. These cells exhibit cell-autonomous morphological changes that are indicative of a basal skew towards an activated phenotype and exaggerated pro-inflammatory cytokine/chemokine release upon immunological challenge. Additionally, AD-derived astrocytes showed elevated Aβ_42_ uptake. The results presented here suggest that PSEN2 (N141I)-mutant glia present with a ‘primed’ phenotype whereby the exposure of Aβ_42_ results in an exaggerated pro-inflammatory response. Whether the increased phagocytic capacity and elevated pro-inflammatory cytokine release from Aβ_42_-stimulated astrocytes and microglia-like cells is detrimental to AD progression, or is initially neuroprotective but gains neurotoxic function as the disease progresses, is not known.

## Abbreviations

AD: Alzheimer’s Disease
fAD: Familial Alzheimer’s Disease
sAD: Sporadic Alzheimer’s Disease
TREM2: Triggering Receptor Expressed On Myeloid Cells 2
APOE: Apolipoprotein
APP: Amyloid Precursor Protein
PSEN1: Presenilin 1
PSEN2: Presenilin 2
Aβ: Amyloid-beta
GFAP: Glial Fibrillary Acidic Protein
iPSC: Induced Pluripotent Stem Cell
TNF-α: Tumor Necrosis Factor-alpha
NPC: Neural Progenitor Cell
FGF2: Fibroblast Growth Factor 2
PBS: Phosphate Buffer Saline
NIM: Neural Induction Medium
DMEM: Dulbecco’s Modified Eagles Medium
HPC: Hematopoietic Progenitor Cell
LPS: Lipopolysaccharide
CXCL1: Chemokine (C-X-C Motif) Ligand 1
MIP: Macrophage Inflammatory Protein 1
MIF: Macrophage Migration Inhibitory Factor
ICAM-1: Intracellular Adhesion Molecule 1
BBB: Blood Brain Barrier

## Declarations

### Ethics approval and consent to participate

Approval for the use of iPSCs was gained from the University of Sydney Institutional Biosafety Committee and Human Research Ethics Committee.

### Consent for publication

Not applicable

### Availability of data and materials

The datasets used during the current study are available from the corresponding authors on reasonable request.

### Competing interests

The authors declare that they have no competing interests.

### Funding

The authors’ research is supported by a National Health and Medical Research Council of Australia (NHMRC) Program Grant (APP1132524). M.K is an NHMRC Principal Research Fellow (APP1154692).

### Authors’ contributions

M.A.S contributed to the design of the work, acquisition of data, analysis of data, interpretation of data and drafted the manuscript. S.D.L, G.G.N and C.M contributed to the acquisition of APOE genotyping data and revised the work. S.R.B and M.S purified the amyloid and revised the work. E.L.W conceived the work, contributed to the design of the work, contributed to the cell line acquisition and derivations, the interpretation of data and revised the work. M.K contributed to the conception of the work and revised the work. All authors have read and approved the submitted version.

## Acknowledgements

The authors acknowledge the technical and scientific assistance of Sydney Microscopy & Microanalysis, the University of Sydney node of Microscopy Australia. We also would like to acknowledge the NYSCF and Cedars-Sinai Medical Center’s David and Janet Polak Foundation Stem Cell Core Laboratory providing the iPSC lines used within the study.

## Notes

### Competing Interest Statement

The authors have declared no competing interest.

