## Supplementary Information File for "iPSC-Derived PSEN2 (N141I) Astrocytes and Microglia Exhibit a Primed Inflammatory Phenotype"

**Additional File 1**

**Table S1** Primary antibodies used for immunofluorescence

| **Target** | **Antibody Species** | **Vendor** | **Product Number** | **Dilution** |
| --- | --- | --- | --- | --- |
| Nestin | mouse | Stem Cell | 60091 | 1:2000 |
| Pax6 | rabbit | Abcam | ab5790 | 1:50 |
| Oct3 | mouse | Stem Cell | 60093.1 | 1:1000 |
| GFAP | rabbit | Abcam | ab7260 | 1:500 |
| S100β | mouse | Sigma Aldrich | S2532 | 1:1000 |
| Iba1 | rabbit | Wako | 019-19741 | 1:500 |
| TREM2 | goat | R&D Systems | AF1828 | 1:100 |
| CX3CR1 | rabbit | Biorad | AHP1589 | 1:250 |

**Table S2** Secondary antibodies used for immunofluorescence

| **Species Reactivity** | **Host** | **Conjugate** | **Vendor** | **Product Number** | **Dilution** |
| --- | --- | --- | --- | --- | --- |
| Mouse | Donkey | Alexa Fluor 488 | ThermoFisher | A-21202 | 1:200 |
| Rabbit | Donkey | Alexa Fluor 594 | ThermoFisher | A-21207 | 1:200 |
| Goat | Donkey | Alexa Fluor 488 | ThermoFisher | A-11055 | 1:200 |

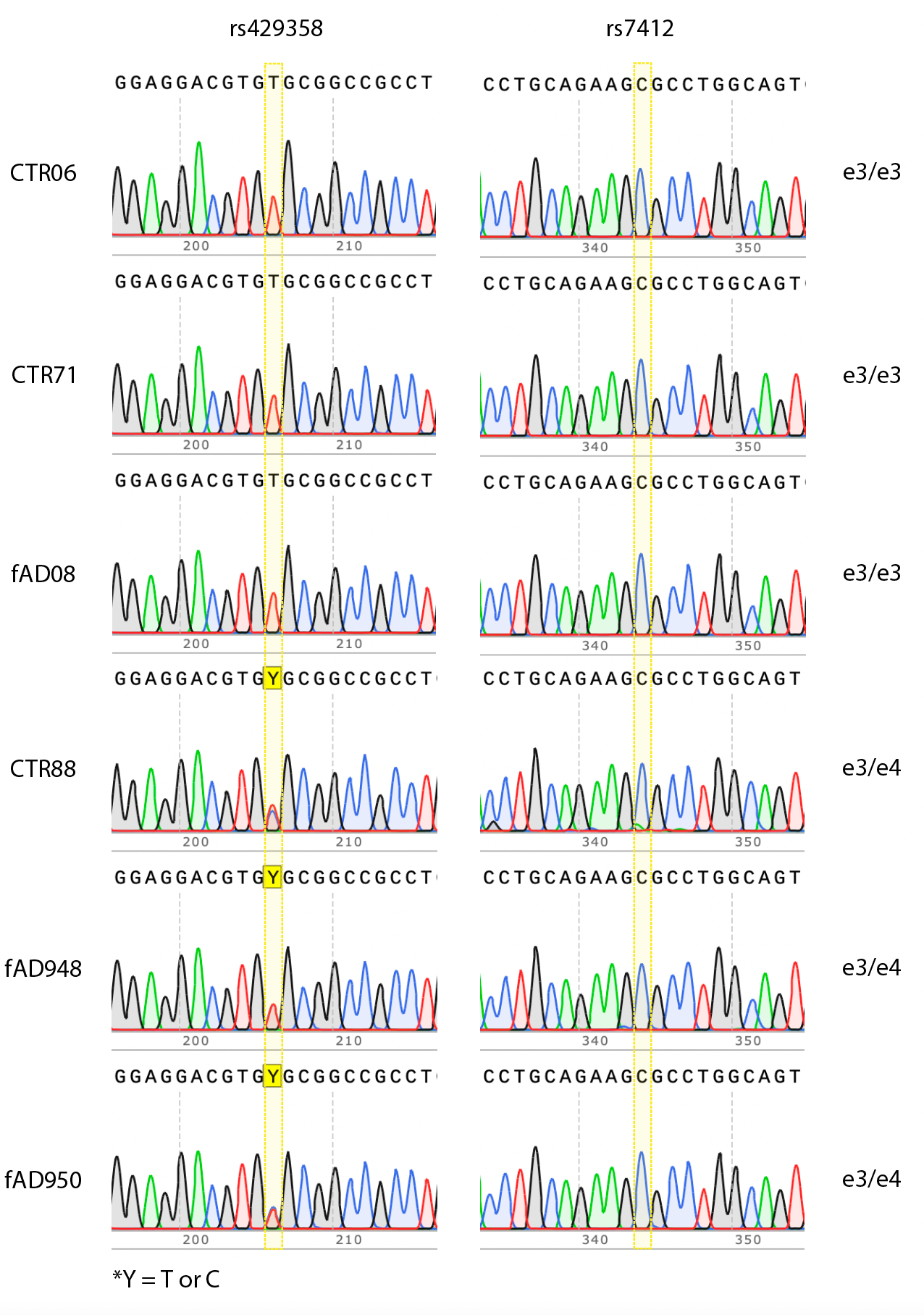

**Fig. S1** Sanger sequencing chromatograms showing APOE genotyping of codon 112 (rs429358) and codon 158 (rs7412) for all iPSC lines. Yellow highlight indicates the position of the single nucleotide polymorphism.

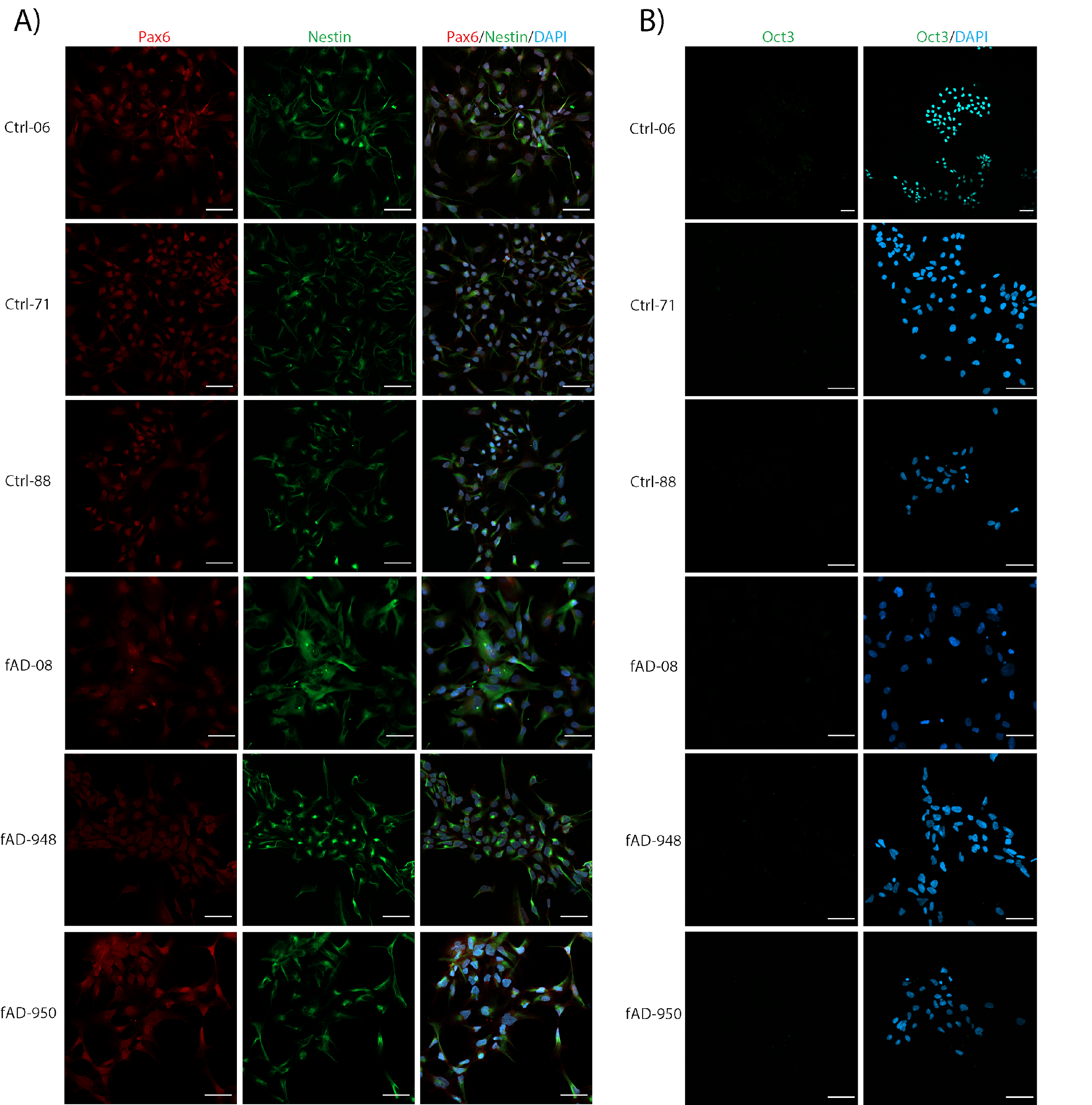

**Fig. S2** Immunofluorescence images of iPSC-derived NPCs from three healthy control lines (Ctrl-06, Ctrl-71, Ctrl-88) and three familial AD lines harbouring a PSEN2 (N141I) mutation (fAD-08, fAD-948, fAD-950). The cells were stained for A) the neural progenitor markers Pax-6 (red) and Nestin (green), B) a pluripotency marker Oct3 (green) and all nuclei were counterstained with DAPI (blue). Scale bars = 50 μm.

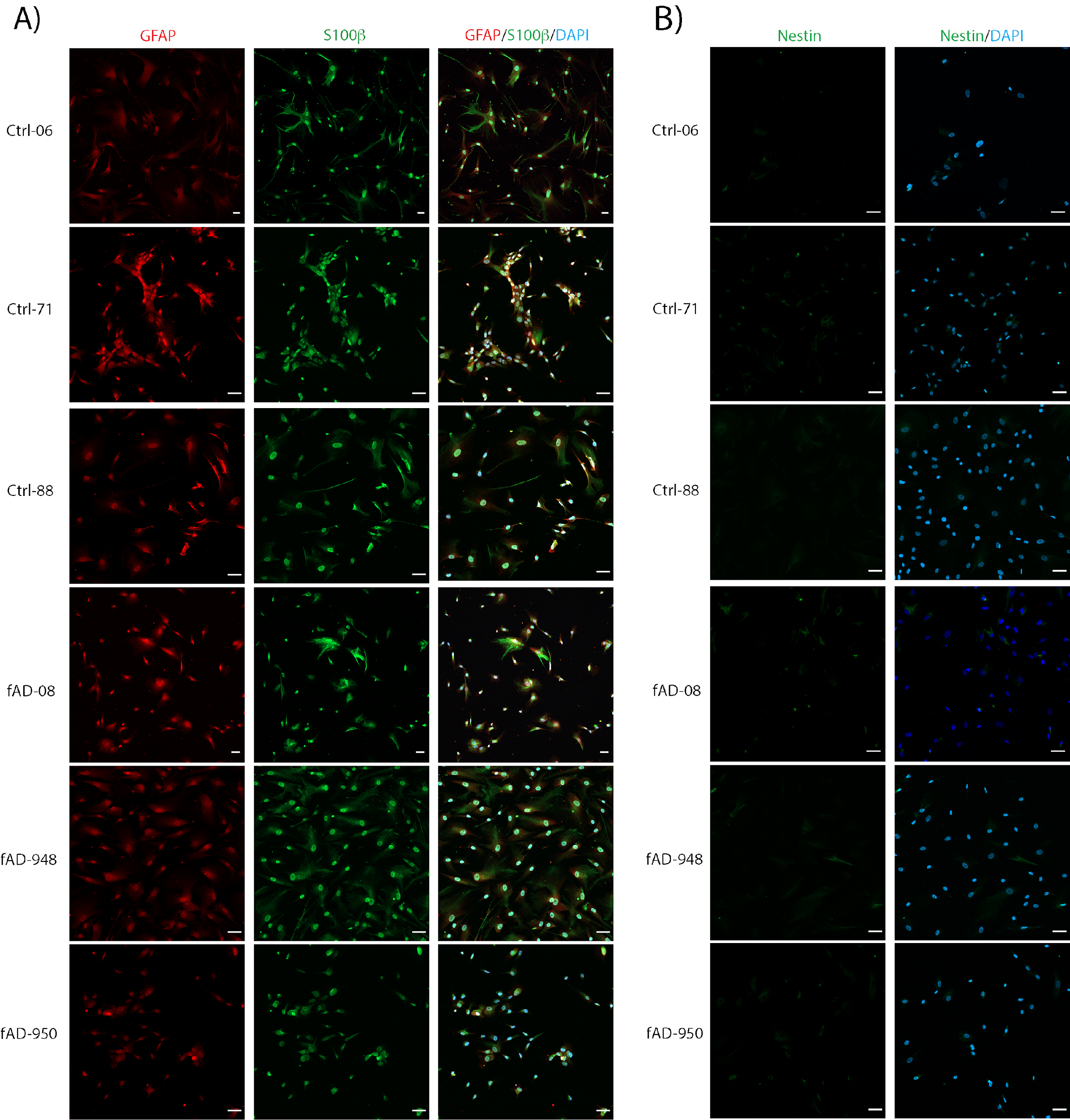
 **Fig. S3** Immunofluorescence images of iPSC-derived astrocytes from three healthy control lines (Ctrl-06, Ctrl-71, Ctrl-88) and three familial AD lines harbouring a PSEN2 (N141I) mutation (fAD-08, fAD-948, fAD-950). The cells were stained for A) astrocyte markers GFAP (red) and S100β (green), B) the NPC marker nestin (green). All nuclei were counterstained with DAPI (blue). Scale bars = 50 μm.

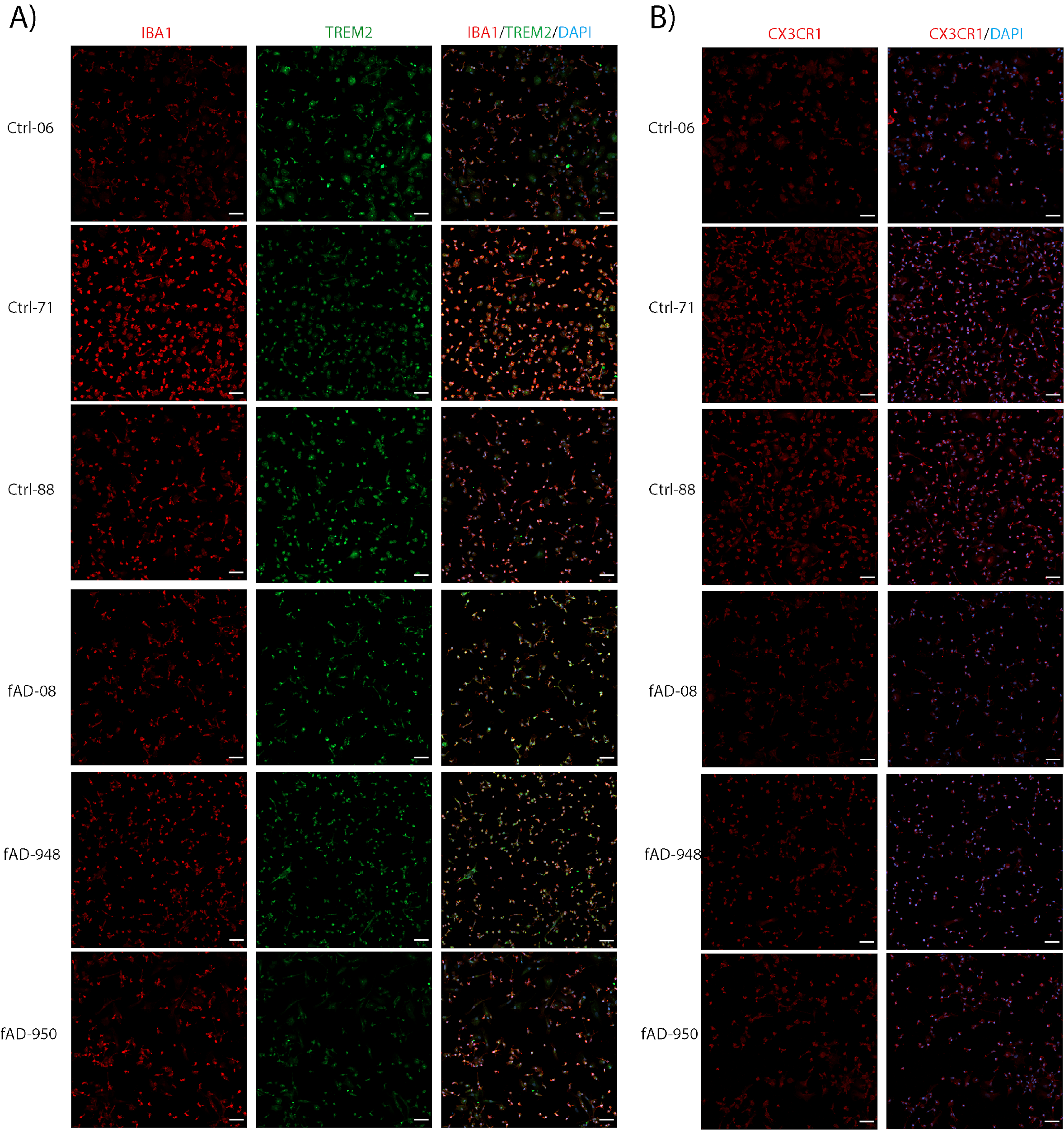

**Fig. S4** Immunofluorescence images of iPSC-derived microglia-like cells from three healthy control lines (Ctrl-06, Ctrl-71, Ctrl-88) and three familial AD lines harbouring a PSEN2 (N141I) mutation (fAD-08, fAD-948, fAD-950). Images show cells stained for A) the microglial markers IBA1 (red), TREM2 (green), B) CX3CR1 (red) and all nuclei were counterstained with DAPI (blue). Scale bars = 50 μm.

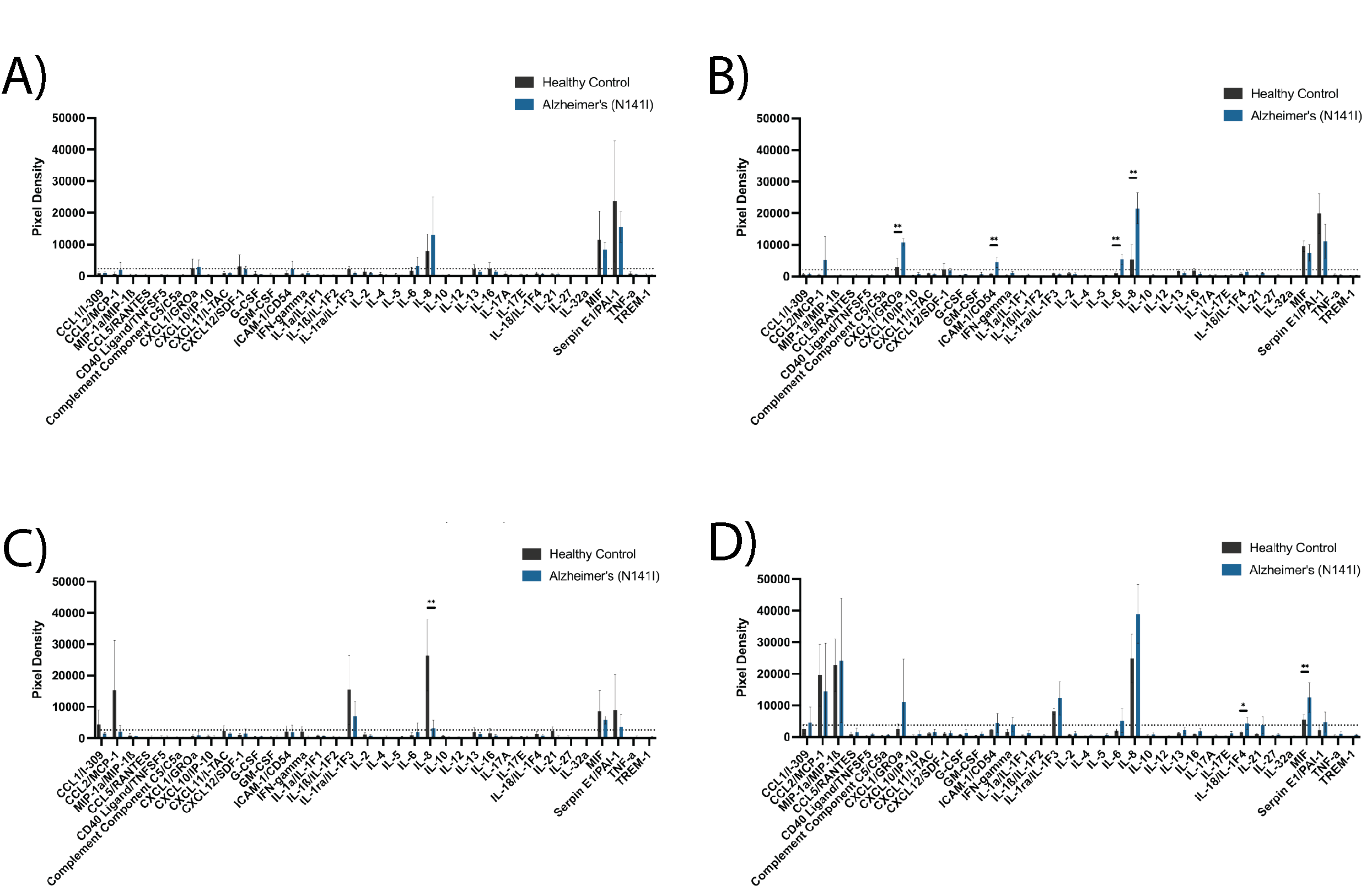

**Fig. S5** Multi-cytokine array of Alzheimer’s or healthy iPSC-derived astrocytes A) basally and B) after 24 h exposure to 10 μM Aβ_42_ and iPSC-derived microglia-like cells C) basally and D) after 24 h exposure to 10 μM Aβ_42_. The figure displays the mean ± SD of three cell lines with the average of two experimental duplicates per line. Multiple unpaired, non-parametric Mann-Whitney t-tests adjusting for a 0.05 false discovery rate were used to test whether there were statistically significant differences between mean cytokine/chemokine release of AD-derived and healthy control astrocytes and microglia-like cells (* p < 0.05, ** p < 0.01). Cytokines that yielded an average intensity value less than 10% of the maximum (represented by the dotted line) were considered background and not included in the statistical analysis.

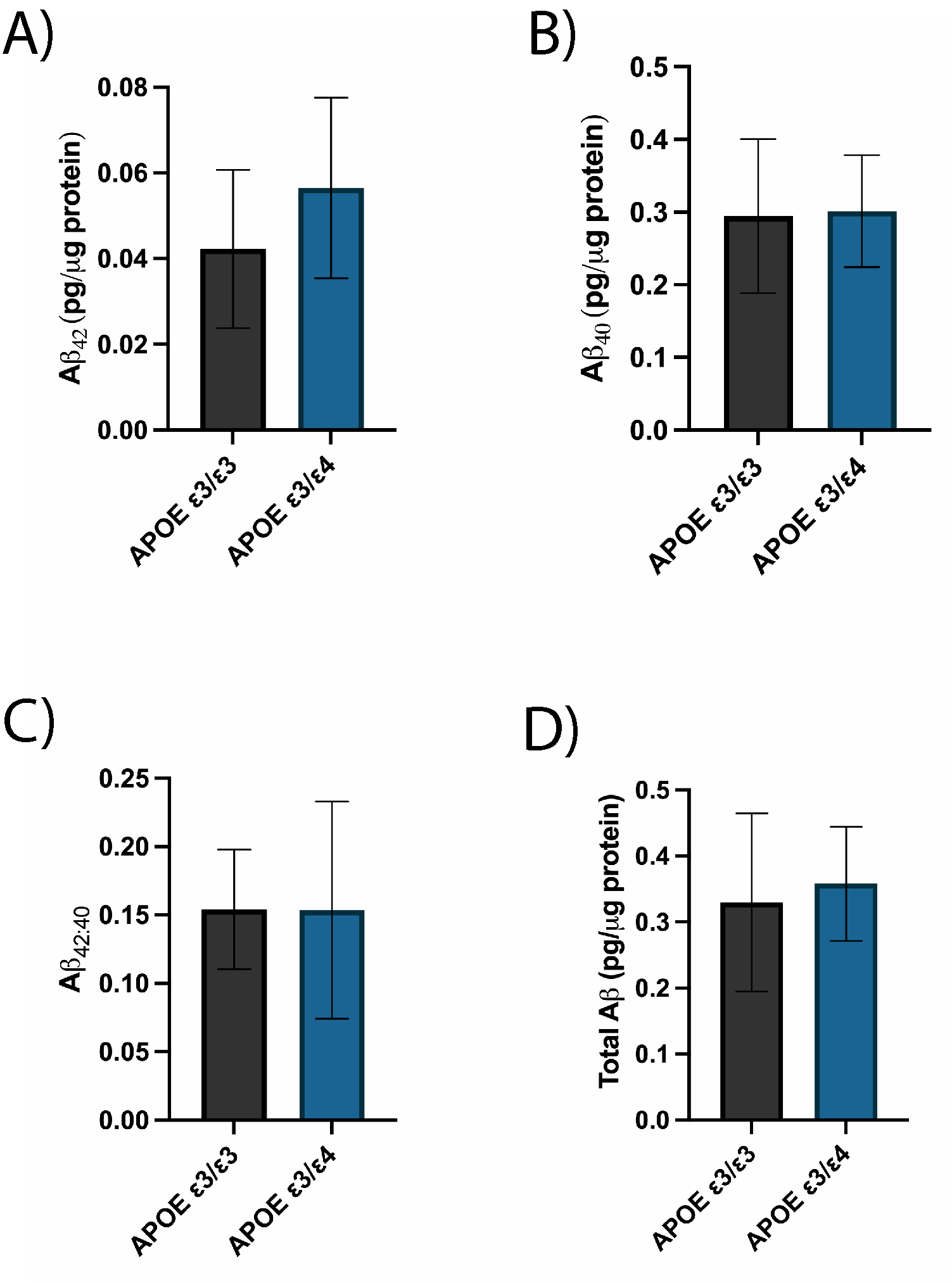

**Fig. S6** Concentrations of A) Aβ_42_, B) Aβ_40_, C) the ratio of Aβ_42:40_ and D) total Aβ protein quantified from iPSC-derived astrocyte supernatants 72 h after plating. Secreted Aβ concentrations were measured using a highly-sensitive ELISA and normalised to total protein concentration determined by BCA. The figure displays the mean ± SD of three cell lines with n ≥ 2 independent experiments per line. A post-hoc unpaired t-test was used to test whether there were statistically significant differences between mean of APOE ε3/ε3 and APOE ε3/ε4 astrocytes.
